# Glycophagy is an ancient bilaterian pathway supporting metabolic adaptation through STBD1 structural evolution

**DOI:** 10.1101/2024.11.07.622431

**Authors:** Liting Ren, Yitian Bai, Chenyu Shi, Ying Tan, Shuyan Zhao, Qi Li, Daniel J Macqueen, Shikai Liu

## Abstract

The selective autophagy of glycogen (glycophagy) has recently emerged as being crucial to glucose homeostasis in vertebrates, yet its origins remain elusive. Here, we provide evidence that starch-binding domain-containing protein 1 (STBD1), the key glycophagy receptor in vertebrates, is functionally conserved in the Pacific oyster, revealing its conserved position within ancient autophagy networks. We show that STBD1 in oysters - as seen in other invertebrate groups - possesses an N-terminal carbohydrate binding module family 20 (CBM20) domain, representing the ancestral state for this protein, while a shuffling of CBM20 to the C-terminus occurred during early chordate evolution. Structural modelling and functional studies reveal that the N-terminal CBM20 organization of STBD1 enhances glycogen binding. Functional experiments demonstrate that an STBD1-glycogen complex, anchored by GABARAPL2, facilitates an increased glycogen flux into autophagosomes for lysosomal degradation. We conclude that glycophagy is deeply conserved in bilaterians and that STBD1 structural evolution underlies potentially adaptive variation in metabolic strategies across distinct animal clades.

## Introduction

All animals rely on conserved metabolic processes to maintain energy balance and cellular homeostasis, essential for survival and fitness in changing environments (*1–3*), including by managing the use of available energy reserves (*4, 5*). Autophagy is an ancient catabolic pathway that maintains cellular homeostasis when energy is limiting (*6*). Unlike canonical metabolic pathways, autophagy operates at low baseline levels under normal conditions, but is activated in response to stressors including starvation (*7, 8*). Autophagy selectively targets intracellular nutrient macromolecules for degradation to supply a source of energy, including glycogen (glycophagy) (*9*) and lipid droplets (lipophagy) (*10*). Importantly, different metazoan clades are thought to primarily rely on distinct energy sources. Vertebrates and insects, for instance, use lipid as a predominant energy store, owing to its high energy density per unit mass (*11–15*). A number of studies have shown that lipophagy is crucial for energy homeostasis in vertebrates (*16–18*), while inhibiting autophagy in *Drosophila* diminished adipocyte differentiation and lipid droplet size without impacting glycogen mobilization (*19*). On the other hand, studies on oysters indicate that glycogen rather than lipid is the primary energy source mobilized for gametogenesis (*20, 21*), strongly associated with an increase of autophagic vesicles (*22*). These findings suggest that nutrient mobilization strategies vary across the Metazoa, and this may be associated with lineage-specific adaptations in metabolism.

Glycophagy is a selective form of autophagy for transporting and degrading glycogen (*23*). In vertebrates, starch-binding domain-containing protein 1 (STBD1) acts as the primary cargo receptor in this process (*24–26*). Under nutrient deprivation, STBD1 recruits glycogen to form STBD1-glycogen complexes, binding with autophagy-related protein 8 (Atg8) family members to assemble glycophagosomes. These glycophagosomes fuse with lysosomes for glycogen degradation, mediated by lysosomal acid α-glucosidase (GAA) (*9*). Mammalian STBD1 contains the hydrophobic N-terminal domain and Atg-interacting motifs (AIMs), as well as a carbohydrate binding module family 20 (CBM20) domain at the C-terminus (*27*). Mutations in CBM20 diminish the stability of STBD1 and its ability to bind glycogen, while attenuating its interactions with glycogen-associated proteins (*28*). Protein structural modelling revealed that CBM20 forms a distorted β-barrel structure with a flexible loop at the C-terminus of human STBD1 (*29*). Interestingly, experimentally relocating CBM20 to the N-terminus of STBD1 significantly enhanced its starch binding and degradation capabilities (*30*). Work to date has failed to reveal the evolutionary origins of glycophagy, including its key cargo receptor STBD1, or whether this autophagy pathway has impacted metabolic adaptations across different animal groups. Bivalve oysters exhibit distinctive glycogen metabolism strategies compared to most metazoans (*31–33*). For instance, glycogen content in oysters is around 7-13-fold higher that of human, fish, and *Drosophila* (fig. S1), while glycogen mobilization is highly regulated by the environment and both nutritional and reproductive status, which is not the case for lipid or protein stores (*33–36*). As oysters have evolved to use glycogen metabolism as a primary strategy for energy homeostasis during stressful conditions, they provide an important system to explore potential adaptations in glycophagy.

In this study, we establish a key role for glycophagy in metabolic regulation in the Pacific oyster, *Magallana gigas*. STBD1 is present in oysters, but has an N-terminal CBM20 domain, and stably binds and degrades glycogen with high efficiency. N-terminal CBM20 represents the ancestral STBD1 organization, arising in the common ancestor of bilaterians and currently encoded in a range of lophotrochozoan genomes. We propose that STBD1-mediated glycophagy is an ancestral strategy for maintaining cellular homeostasis during stress, which likely supported metabolic adaptations during bilaterian evolution.

## Results

### STBD1-mediated glycophagy plays a crucial role in energy metabolism in oysters

To investigate the role of autophagy in the response of oysters to stress, we manipulated nutritional status in *M. gigas*, using a 14-day fasting period followed by a refeeding phase (Fig. 1A). Immunohistochemistry showed that the autophagy marker LC3 was significantly enhanced by fasting, with signal weakening after refeeding (Fig. 1B and fig. S2). In contrast, glycogen signals markedly diminished after fasting and subsequently reaccumulated upon refeeding, whereas lipid signals exhibited no changes (Fig. 1B, figs. S2 and S3). Notably, LC3 signal was concentrated in the ciliated epithelium (CE) and vesicular connective tissue (VCT) cells of labial palp, the adult tissue with the highest glycogen content (fig. S4). The high correlation between LC3 and glycogen signals, and the low correlation with lipid signals (Fig. 1B), implicates a specific role for autophagy in glycogen depletion under nutrient stress.

**Fig. 1.**
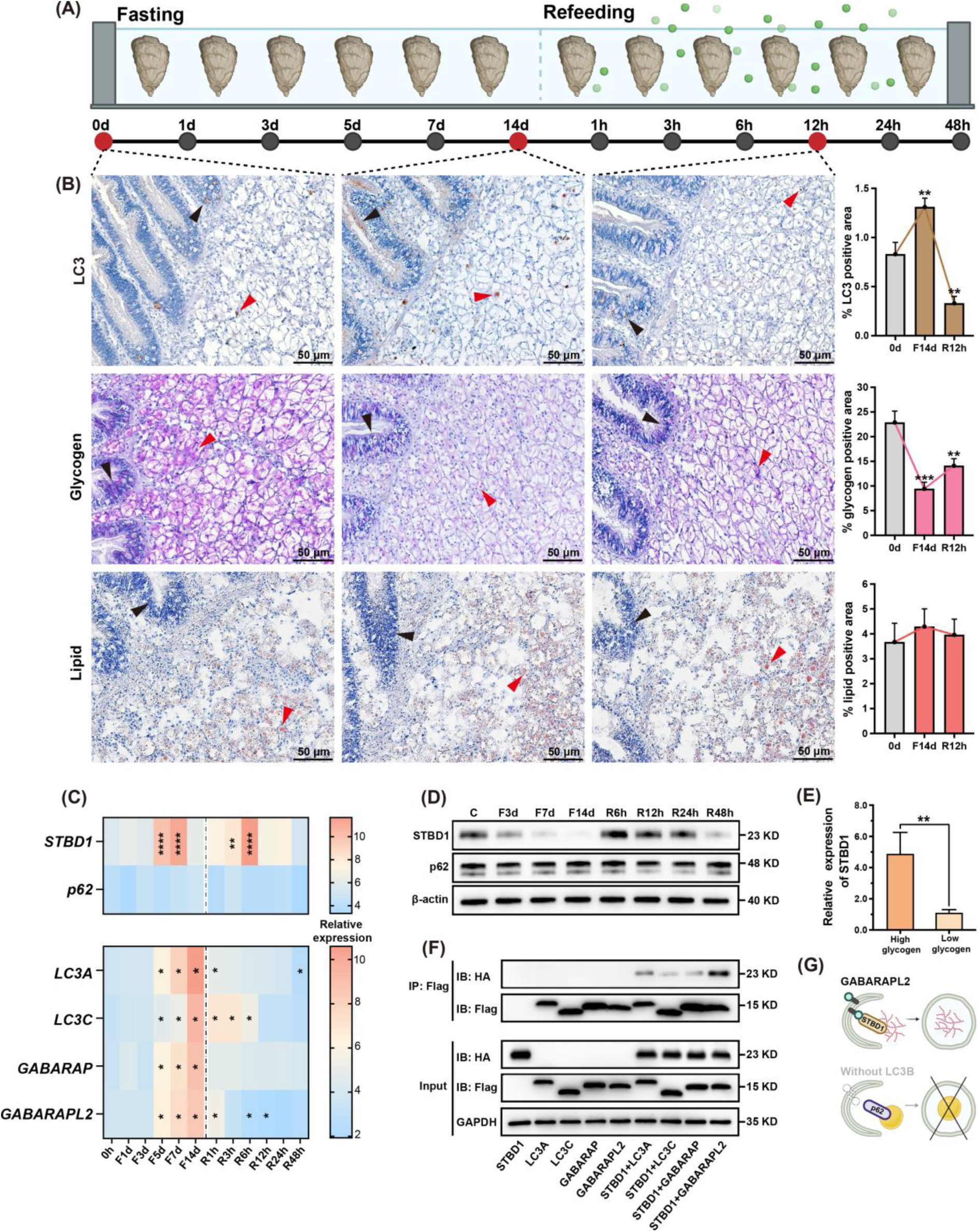
STBD1-mediated glycophagy, rather than p62-mediated lipophagy, serves as a primary energy source in oysters. **(A)** Nutritional status in *M. gigas* was manipulated through a 14-day (d) period of fasting followed by a refeeding period lasting 48 hours (h). **(B)** Signals for the autophagy marker LC3 (brown), glycogen (pink), lipid (red) in labial palp were observed after 14 d of fasting followed by 12 h of refeeding. Arrows indicate dense distribution of signals in the CE (black) and VCT (red) cells within the labial palp. Scale bars represent 50 μm. Positive signals was quantified using ImageJ, as displayed in graphs on the right (\*\**P* < 0.01, \*\*\**P* < 0.001). **(C)** Expression levels of autophagy-related genes, *STBD1*, *p62*, *LC3A*, *LC3C*, *GABARAP* and *GABARAPL2*, in the labial palp of oysters after 14 d fasting followed by 48 h refeeding (*N* = 6, mean ± SD, \**P* < 0.05, \*\**P* < 0.01, \*\*\*\**P* < 0.0001). (**D)** Protein levels of STBD1 and p62 in the labial palp after 14 d fasting followed by 48 h refeeding. **(E)** Expression levels of *STBD1* in oyster individuals showing high and low glycogen content (*N* = 9, mean ± SD. \*\**P* < 0.01). **(F)** Interaction between STBD1 and either LC3A, LC3C, GABARAP or GABARAPL2 using co-immunoprecipitation assays. Western blotting detected bands from STBD1 (HA-conjugated) and LC3A, LC3C, GABARAP and GABARAPL2 (Flag- conjugated). **(G)** Schematic model of glycophagy in oysters: STBD1 interacts with GABARAPL2 directing glycogen into the autophagic degradation pathway, while p62- mediated lipophagy appears absent, possibly due to loss of LC3B. Abbreviations used: F, Fasting; R, Refeeding; d, day; h, hour.

To provide further evidence for the role of autophagy, we assessed the expression of candidate *M. gigas* orthologs for genes encoding the glycophagy receptor STBD1 (LOC105333128) and the lipophagy receptor p62 (LOC105345634) (*37*). *STBD1* mRNA expression increased significantly following 5-days fasting, peaked after 7 days, then declined, with a rapid surge following refeeding, subsequently returning to control levels (Fig. 1C and fig. S5). In contrast, *p62* exhibited no significant alterations in expression across the same experiment. Consistently, STBD1 protein levels gradually decreased during fasting and significantly accumulated after refeeding, while p62 protein levels showed no statistically significant differences (Fig. 1D and fig. S6). These findings indicate that STBD1, rather than p62, plays a primary role in the regulation of autophagy-related energy metabolism in oysters. Moreover, *STBD1* showed increased mRNA expression in oysters with high levels of glycogen content (Fig. 1E and fig S7), suggesting a close association between STBD1 and glycogen metabolism, implicating glycophagy as a source of energy during nutritional deprivation.

The STBD1-glycogen complex binds to the Atg8 family of autophagy-modifying proteins through the AIM motif of STBD1, serving as another essential requirement for promoting the autophagic sequestration of glycogen (*27, 38, 39*). GABARAPL1, an Atg8 family member, interacts with the STBD1-glycogen complex to trigger autophagosome maturation in vertebrates (*24, 27*). We screened the *M. gigas* genome for all known Atg8 family members to establish if GABARAPL1 is conserved in oysters. Phylogenetic analysis (fig. S8) revealed *M. gigas* orthologs for LC3A, LC3C, GABARAP and GABARAPL2, but not GABARAPL1, nor LC3B, an Atg8 family member involved in lipophagy through its interaction with p62 (*16, 17*). In light of their considerable homology, it is possible that the primary Atg family members interacting with STBD1 in *M. gigas* are substituted; mirroring findings in *C. elegans*, which lacks the LC3 homolog LGG-2, with autophagosome formation instead critically dependent on the GABARAP homolog LGG-1 (*40, 41*). The expression of the four identified oyster Atg8 family members increased significantly during fasting and decreased upon refeeding (Fig. 1C). We also identified a clear mRNA co-localization signal for *STBD1* with Atg8 family members in the CE and VCT cells of labial palp, which was strongest for *GABARAPL2*, followed by *GABARAPL* and *LC3A*, while no signal was detected with *LC3C* in the VCT cells (fig. S9). To identify physical interactions between *M. gigas* STBD1 and Atg8 family members, each protein was co-expressed and their binding interactions assessed using co-immunoprecipitation assays (Fig. 1F). The strongest interaction was between STBD1 and GABARAPL2, with an intensity 4.36-fold higher than between STBD1 and the other three Atg8 family members (fig. S10). Therefore, GABARAPL2 is the main interacting partner of STBD1 in oysters, which likely allows the entry of STBD1-glycogen complexes into autophagosomes (Fig. 1G). The potential loss of LC3B in oysters may also inhibit the autophagic degradation of lipid droplets in oysters. Overall, these results support that STBD1-mediated glycophagy is a crucial strategy used by oysters to maintain homeostasis during nutritional deficiency, whereas there is little evidence for an equivalent role for lipophagy.

### N-terminal CBM20 STBD1 domain has higher predicted affinity for glycogen

Given the established role of conserved protein domains in STBD1-mediated glycogen binding and autophagosome degradation, we compared the protein structure of *M. gigas* STBD1 with its orthologs in representative vertebrates *Mus musculus* and *Danio rerio* (Fig. 2, A-C). The most striking difference between *M. gigas* STBD1 and vertebrate STBD1 was that the CBM20 domain was positioned at the N- and C-terminus, respectively. Moreover, the N-terminal hydrophobic domain of vertebrate STBD1, which dictates STBD1 subcellular localization, but is dispensable for glycogenolytic activity (*26, 28*), was absent in *M. gigas*. Potential AIM motifs, which bind Atg8 family members, were identified in *M. gigas* STBD1 (Y110-L113, Y145-V148, and F177-I180). However, characteristic WXXV AIM motif found in vertebrate STBD1 proteins was not present in *M. gigas* STBD1. This discrepancy could be ascribed to the selectivity and preference of Atg8 members for specific AIM types (*42, 43*).

**Fig. 2.**
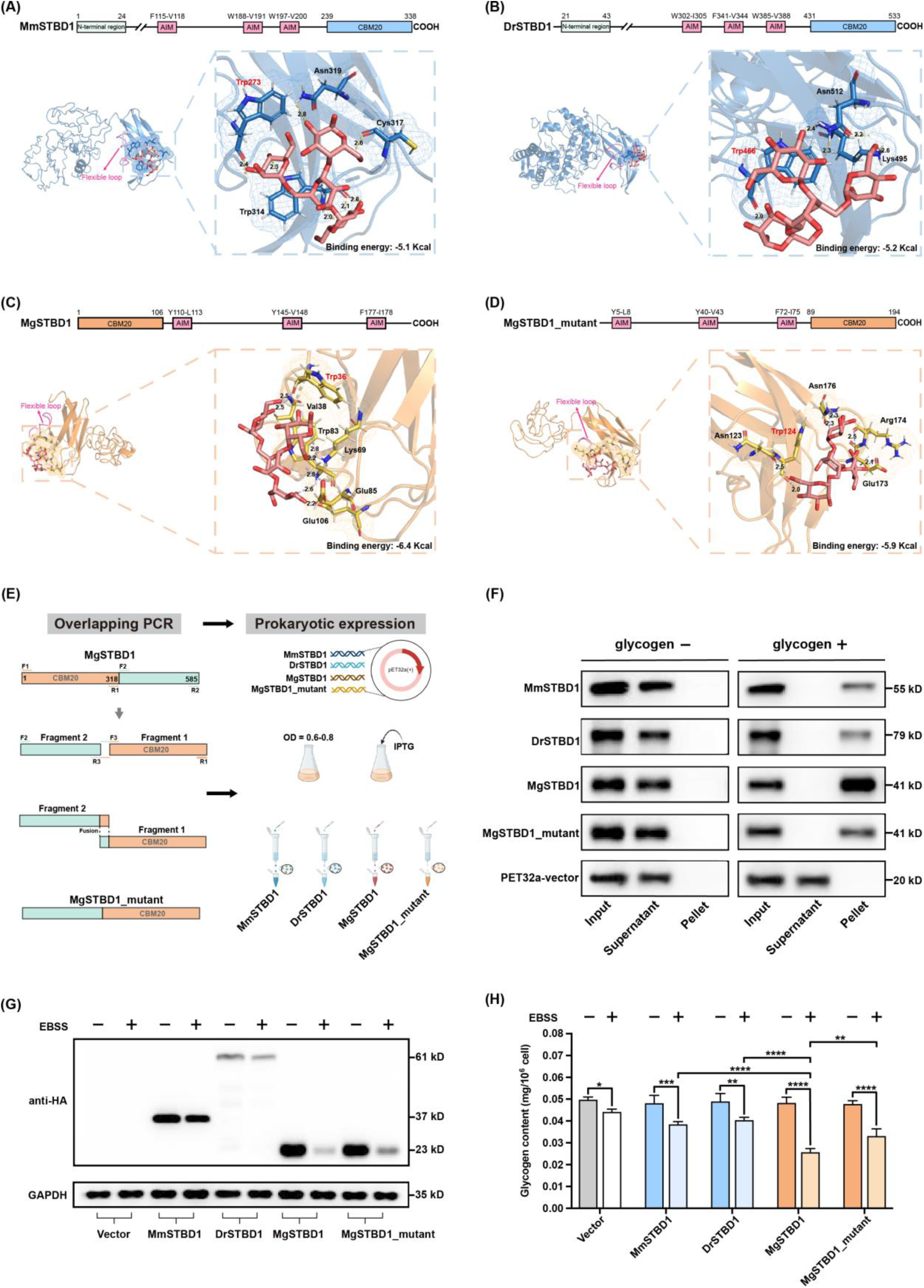
The N-terminal CBM20 domain of oyster STBD1 influences glycogen binding and degradation. (A-D) Comparative analysis of protein structures and interactions of glycogen with mouse, zebrafish and wild-type *M. gigas* STBD1, and an *in silico*-mutated *M. gigas* STBD1 with CBM20 moved to the C-terminus (90-194 aa). Optimal docking models (RMSD < 2) were constructed and binding energies were calculated. Flexible loop structures are indicated by pink arrows. Conserved Trp residues known to form direct hydrogen bonds with carbohydrates are indicated as red letters. Hydrogen bonds are indicated by yellow or purple dashed lines and labeled with bond lengths. **(E)** Schematic diagram summarizing the construction and isolation of a *M. gigas* STBD1 protein with CBM20 moved to the C-terminus, which was overexpressed along with mouse, zebrafish and wild-type *M. gigas* STBD1. **(F)** Binding of the four recombinant STBD1 proteins with glycogen was measured. Western blotting assessed the protein levels in inputs, supernatants, and the pellet after ultracentrifugation. PET32a vector protein was used as a negative control. **(G)** Changes in protein levels of each STBD1 protein overexpressed in 293T cells after transfection for 24 h, followed by treatment with or without EBSS for 6 h. Western blotting was used to verify protein levels. (H) Glycogen content in 293T cells after overexpression of each STBD1 protein for 24 h, followed by treatment with or without EBSS for 6 h (*N* = 4, mean ± SD. \**P* < 0.05, \*\**P* < 0.01, \*\*\**P* < 0.001, \*\*\*\**P* < 0.0001). Abbreviations used: *Mm*, *Mus musculus*; *Dr*, *Danio rerio*; *Mg*, *Magallana gigas*.

To elucidate the impact of the CBM20 domain’s location on STBD1 function, we predicted and compared three-dimensional structures of *M. gigas* STBD1 (N-terminus CBM20, wild-type), vertebrate STBD1 proteins (C-terminus CBM20) and an *in silico* re-arranged *M. gigas* STBD1 (C-terminus CBM20) (Fig. 2D), before predicting interactions of these molecules with glycogen. For all STBD1 proteins, CBM20 was modelled as an open-side and distorted β-barrel, although the twist of β-barrel in *M. gigas* STBD1 was opposite to other STBD1 proteins (Fig. 2, C-D). Molecular docking revealed that the glycogen binding groove was located within a flexible loop, with the hydrogen bonding interaction between the main chain carbonyl oxygen of the conserved Trp in CBM20 and glycogen (*44*). Notably, the binding free energy of *M. gigas* STBD1 with glycogen was -6.4 Kcal/mol, higher than that for all three C-terminus CBM20 proteins (Fig. 2, A-D), which could be attributed to a greater number of residues and hydrogen bonds (fig. S11). Specifically, glycogen was predicted to interact with *M. gigas* STBD1 via seven hydrogen bonds involving Trp36, Val38, Trp83, Glu85 and Glu106, and hydrophobic interactions with Val38, Phe81 and Arg82 (Fig. 2C and fig. S11C). Notably, Trp83 and Lys69 of wild-type *M. gigas* STBD1 were predicted to form hydrogen bonds with glycogen, whereas the corresponding Trp171 and Lys157 in the *in silico* mutant CBM20 C-terminus STBD1 interacted with glycogen via hydrophobic forces (fig. S11D). These structural models support that the N-terminal position of CBM20 enhances the interaction of STBD1 with glycogen, which is predicted to result in higher glycogen binding affinity.

### Oyster STBD1 possesses enhanced binding affinity that expedites glycogen breakdown

Next, we sought to experimentally validate the predicted enhanced glycogen-binding affinity conferred to oyster STBD1 by its distinctive N-terminal CBM20. We successfully isolated recombinant proteins for wild-type STBD1 proteins in zebrafish, mouse and *M. gigas*, along with a mutant *M. gigas* STBD1 protein with CBM20 moved to the C-terminus (Fig. 2E and fig. S12). Glycogen co-sedimentation assays showed that all isolated STBD1 proteins bound glycogen *in vitro*, with the strongest binding for wild-type *M. gigas* (Fig. 2F), approximately 1.6-fold higher than mutant *M. gigas* STBD1, and 2.1-2.7-fold higher than the wild-type vertebrate STBD1 proteins (fig. S13). Thus, the N-terminal position of CBM20 in oyster STBD1 markedly enhances affinity for glycogen compared to both the same protein, as well as evolutionarily distant STBD1 orthologs, containing a C-terminal CBM20 domain.

We next tested the hypothesis that the relatively stronger affinity between glycogen and STBD1 from *M. gigas* supports enhanced glycogen degradation. The same four STBD1 proteins were overexpressed in human 293T cells and treated with the autophagy inducer EBSS for 6 h, before changes in STBD1 protein abundance and glycogen content were determined by comparison to controls. Western blot analysis showed that *M. gigas* STBD1 protein abundance was markedly higher than for vertebrate STBD1 (Fig. 2G). Moreover, a 5.48-fold reduction in wild-type *M. gigas* STBD1 abundance was observed following 6 h autophagy induction, compared to 1.31-fold, 1.91-fold and 2.65-fold for mouse, zebrafish and the mutant *M. gigas* STBD1 (fig. S14), indicating that *M. gigas* STBD1 protein is utilized more efficiently than the three STBD1 proteins with C-terminal CBM20 domains. Importantly, the glycogen content of 293T cells overexpressing wild-type *M. gigas* STBD1 was significantly lower after autophagy induction than for the other three STBD1 proteins (Fig. 2H). Nonetheless, 293T cells overexpressing all four STBD1 proteins showed significant reductions in glycogen content following autophagy induction compared to controls (Fig. 2H). Together, these results indicate that wild-type STBD1 protein with an N-terminal CBM20 domain possesses higher glycogen binding affinity, facilitating a greater influx of glycogen into the autophagy pathway and more rapid glycogen decomposition.

### Resolving the evolutionary origins of STBD1 in metazoans

Our results prove that glycophagy is conserved between oysters and vertebrates, revealing the significance of the CBM20 domain’s position for STBD1 glycogen binding function. Considering the vital role of CBM20 in glycophagy, we investigated the evolutionary dynamics of genes encoding CBM20 domain-containing proteins across metazoan lineages, with the aim of defining the phylogenetic origin of STBD1, including its CBM20 domain arrangement. We conducted a comprehensive genome-wide scan of CBM20 domain containing proteins in 61 species from 13 phyla (table S1) and compared their domain organizations and phylogenetic relationships using a maximum likelihood approach (Figs. 3 and 4).

**Fig. 3.**
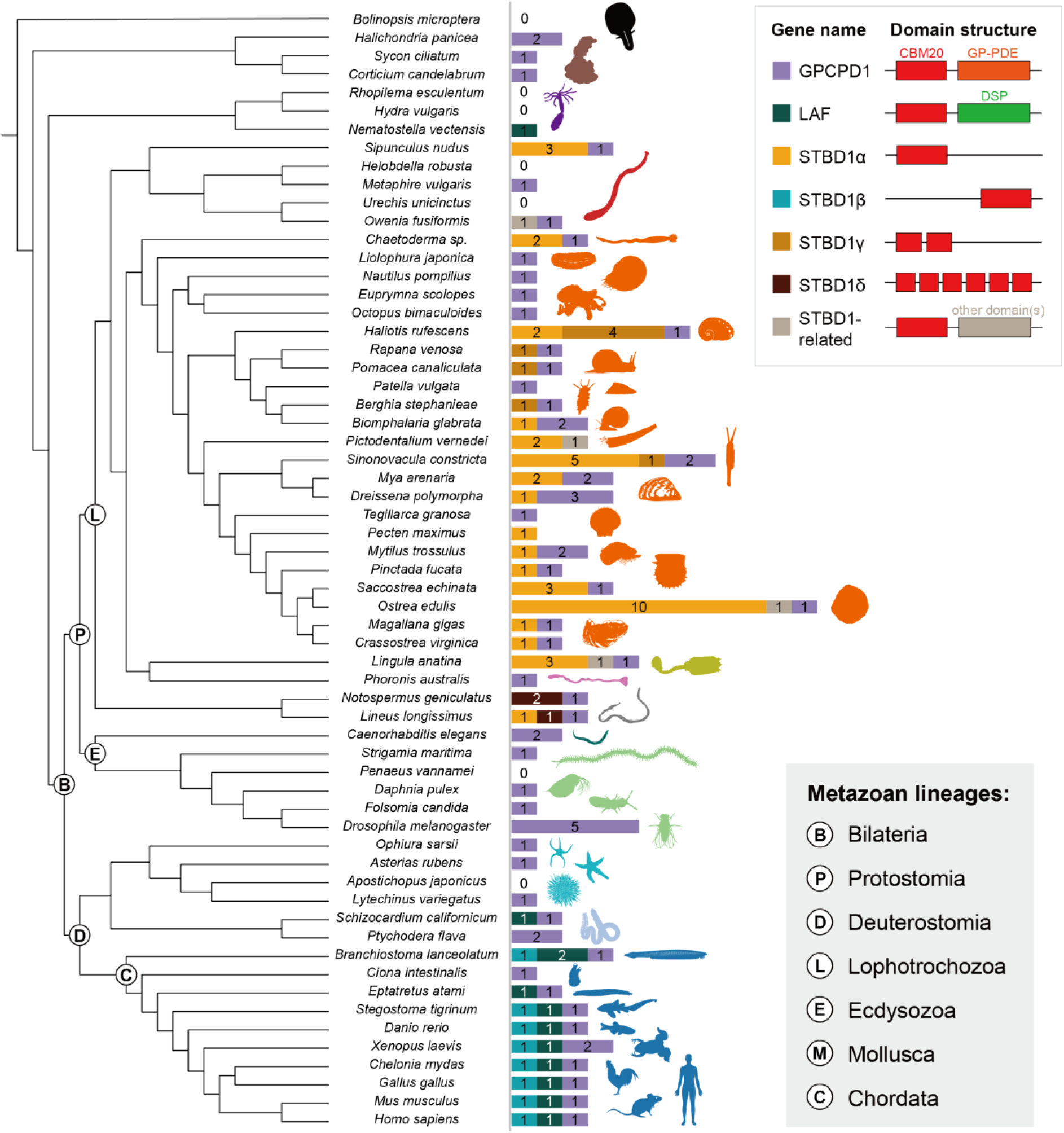
Phylogenetic distribution of genes encoding proteins with CBM20 domains across metazoans. The schematic tree indicates evolutionary relationships based on NCBI Taxonomy. CBM20 domain containing proteins were categorized into seven types based on domain structures (fig. S15), with schematic diagrams in the upper-right corner. Other domains refer to EF-hand (IPR002048), peptidase M12B (IPR001590), IPT (IPR002909) and glycosyl hydrolase family 13 (IPR006047). Abbreviations used: CBM20, Carbohydrate binding module family 20 domain (IPR002044); GP-PDE, Glycerophosphodiester phosphodiesterase domain (IPR030395); DSP, Dual specificity phosphatase domain (IPR000340).

**Fig. 4.**
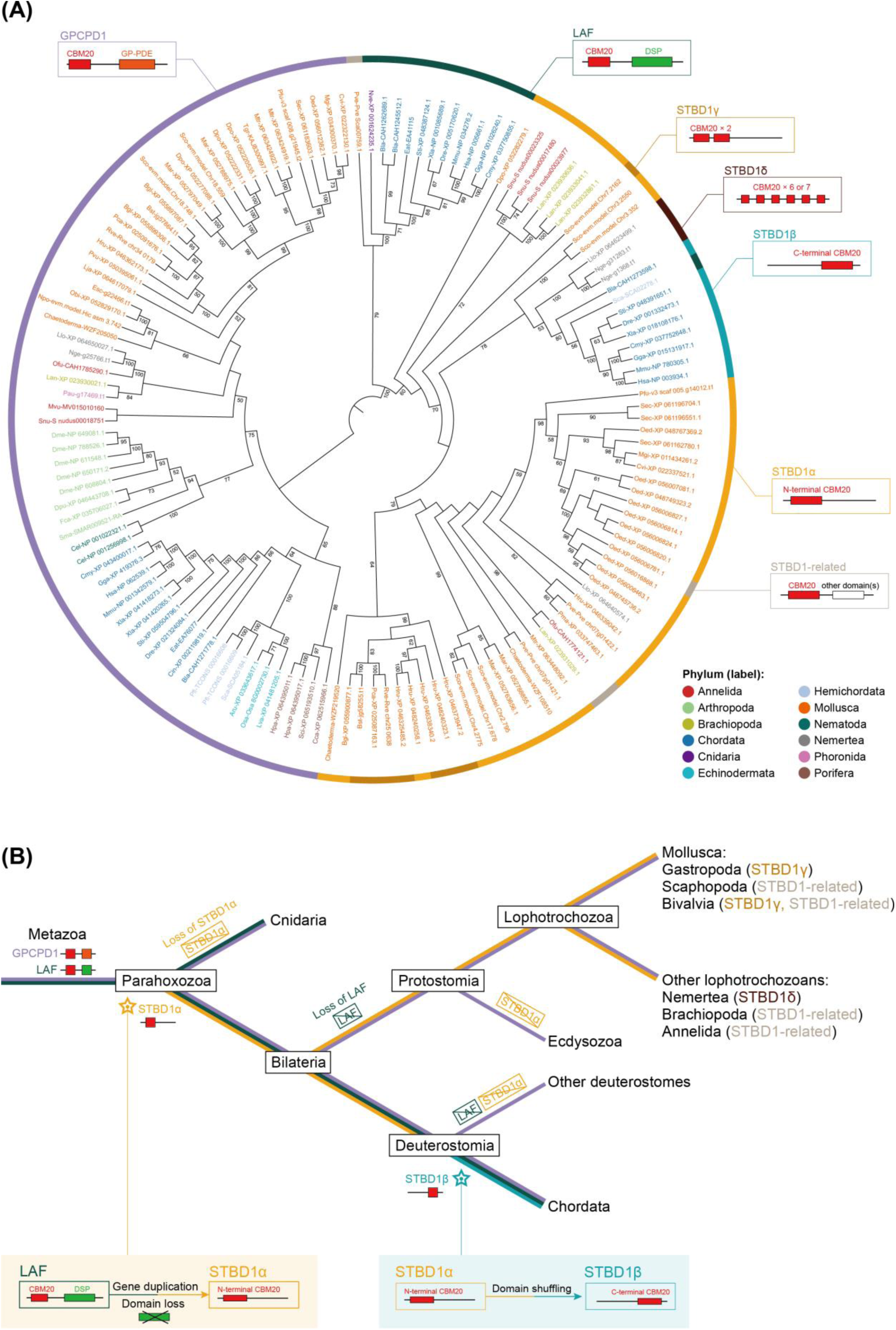
Evolutionary history of CBM20 domain proteins in metazoans. **(A)** Maximum likelihood phylogenetic analysis of metazoan STBD1, LAF, and GPCPD1 proteins. The colors of protein IDs of species represent distinct lineages according to the provided tree. The outer strips (indicating types of domain architecture) are colored following the rule in Figure 3. Numbers shown on the nodes are bootstrap values (> 50%). The corresponding abbreviations of species names are shown in Table S1. **(B)** Hypothesis for STBD1 evolution: the common ancestor of parahoxozoans evolved STBD1α by gene duplication of LAF, potentially followed by loss of the DSP domain. STBD1α was independently lost in Cnidaria, Ecdysozoa and Echinodermata, and evolved into STBD1γ, STBD1δ, or STBD1-related genes in mollusks, nemerteans, annelids, and brachiopods. At the root of Chordata, the STBD1β gene evolved from STBD1α through domain shuffling. Protostomes and echinoderms independently lost LAF. The yellow, blue, dark green, and purple branches indicate the presence of STBD1α, STBD1β, LAF, and GPCPD1 genes during evolution, respectively.

Proteins containing CBM20 were present in all examined phyla except Ctenophora, representing seven major domain architectures (Fig. 3 and fig. S15). These proteins form three clades: glycerophosphocholine phosphodiesterase 1 (GPCPD1), laforin (LAF), and STBD1, each including previously characterized vertebrate representatives (Fig. 4A), consistent with previous reports in mammals that CBM20 is restricted to GPCPD1, LAF and STBD1 (*27*). Despite being absent in protostomes and echinoderms, a cnidarian protein within the LAF clade suggests that LAF was present before the divergence of cnidarians and bilaterians. The presence of poriferan sequences in the GPCPD1 clade, alongside the fact that GPCPD1, LAF and STBD1 groupings predate branching events within each of these clades, is consistent with all three CBM20-containing clades being present in the metazoan ancestor, with subsequent lineage-specific patterns of loss and expansion. This includes evidence for prevalent losses of LAF proteins, alongside widespread retention of GPCPD1 in different metazoan lineages (Fig. 4A).

Notably, STBD1 proteins with a single C-terminal CBM20 domain (hereafter: STBD1α) were exclusively found in lophotrochozoans, while N-terminal CBM20 (hereafter: STBD1β) proteins were restricted to Chordata (Figs. 3 and 4A). A marked STBD1α expansion was observed in mollusks, especially the bivalves *Sinonovacula constricta* and *Ostrea edulis*. In addition, we identified STBD1 proteins with two CBM20 domains (STBD1γ) in *S. constricta* and some gastropods, and multiple CBM20 domains (STBD1δ) in nemerteans. Thus, a diverse repertoire of STBD1 proteins have evolved in lophotrochozoans and chordates, implying the potential for functional divergence and specialization.

LAF and STBD1 were supported to represent sister clades using midpoint rooting (*45*) of our phylogenetic tree (Fig. 4A). Given that STBD1 is present only in bilaterians, it is challenging to distinguish loss scenarios in other metazoan lineages from an origin specific to early bilaterian evolution. Nonetheless, a cnidarian LAF sequence suggests that STBD1 and LAF originated no later than the common ancestor of bilaterians and cnidarians. Furthermore, STBD1α exhibits a similar N-terminal CBM20 domain structure to LAF and GPCPD1. Therefore, we propose that STBD1α represents the ancestral domain architecture of STBD1, which originated from LAF through gene duplication and domain loss in the common ancestor of parahoxozoans (Fig. 4B). After the divergence from other deuterostomes, chordate-specific STBD1β may derive from domain shuffling of CBM20 from the N-terminus to the C-terminus. Moreover, lineage-specific domain duplication events in nemerteans and mollusks have resulted in the generation of STBD1γ and STBD1δ with two or more CBM20 domains. In lophotrochozoans, species-specific gene duplications have expanded the STBD1 repertoire. Overall, gene expansion, domain shuffling and duplication have shaped STBD1 evolution to the present various degrees of conservation and divergence observed across bilaterians.

## Discussion

Glycogen, the primary reservoir for surplus glucose, facilitates rapid energy mobilization during periods of energy deficit due to its uncomplicated chemical and molecular mass compared to lipids (*46, 47*). Although canonical glycogen metabolism pathways are highly conserved across metazoans, the efficiency of glycogen storage and utilization varies remarkably among lineages (*12, 48–50*). Glycophagy recently emerged as a non-canonical energy producing pathway in vertebrates (*23–25*). A recent study revealed that oysters undergoing energetically demanding gametogenesis showed an increased number of autophagic vesicles, serving as large reservoirs of glycogen (*22*), suggesting a potential role for glycophagy in energy homeostasis, and hinting that this pathway may considerably predate vertebrates. Using oysters as a study system, this study comprehensively resolves the evolutionary origins of glycophagy within Metazoa, revealing an overall conserved role of STBD1 within ancient autophagy networks, alongside evidence for structural modifications shaping glycogen binding that can be linked to the distinct metabolic strategies of different metazoan lineages.

Protein domain evolution is a key process leading to changes in molecular function within metabolic pathways (*3, 51*). CBM20 represents the archetypal starch-binding domain, originally characterized as located at the C-terminus of bacterial amylolytic enzymes (*52, 53*). In contrast, CBM20 in non-amylolytic enzymes such as the LAF and GPCPD families, is always located at the N-terminus (*44, 54, 55*). Our results demonstrate that CBM20 was also ancestrally located at the N-terminal of STBD1, when it arose during early bilaterian evolution, potentially from duplication of LAF. A domain shuffling event in early chordates led to the derived C-terminal CBM20 of STBD1 found in all vertebrates (Fig. 4B). Conversely, oyster STBD1 exhibits the ancestral N-terminal CBM20 domain, which enhances glycogen binding efficiency according to several lines of evidence in this study. This finding is consistent with a recent study, which engineered glycoside hydrolase to have a N-terminal (instead of wild-type C-terminal) CBM20 domain, leading to enhanced starch degrading activity (*30*). The repositioning of CBM20 to the C-terminus of STBD1 thus led to lower binding with glycogen compared to the ancestral condition. It is plausible this was adaptively linked to the metabolic strategy of vertebrates, where lipophagy serves as a complement to glycophagy (*16, 25*), balancing energy supply and nutrient mobilization; an adaptation evidently lacking in oysters. Interestingly, STBD1 has evolved rapidly in lophotrochozoans, with a high proportion of lineage-specific gene duplications and domain rearrangements (Fig. 4). The rapid diversification of STBD1 may contribute to the adaptive evolution of glycophagy pathways in lophotrochozoans. Further work is required to elucidate the functional role of these diverse STBD1 homologs and determine whether these putative glycophagy receptors also interact with upstream autophagy components.

In conclusion, the conserved function of STBD1 in oysters and vertebrates positions STBD1-mediated glycophagy as an ancestral metabolic pathway in bilaterians (Fig. 5). Our findings further highlight the importance of protein structural evolution in the regulation of energy metabolism, thereby advancing understanding of how distinct metabolic strategies arose in different animal lineages.

**Fig. 5.**
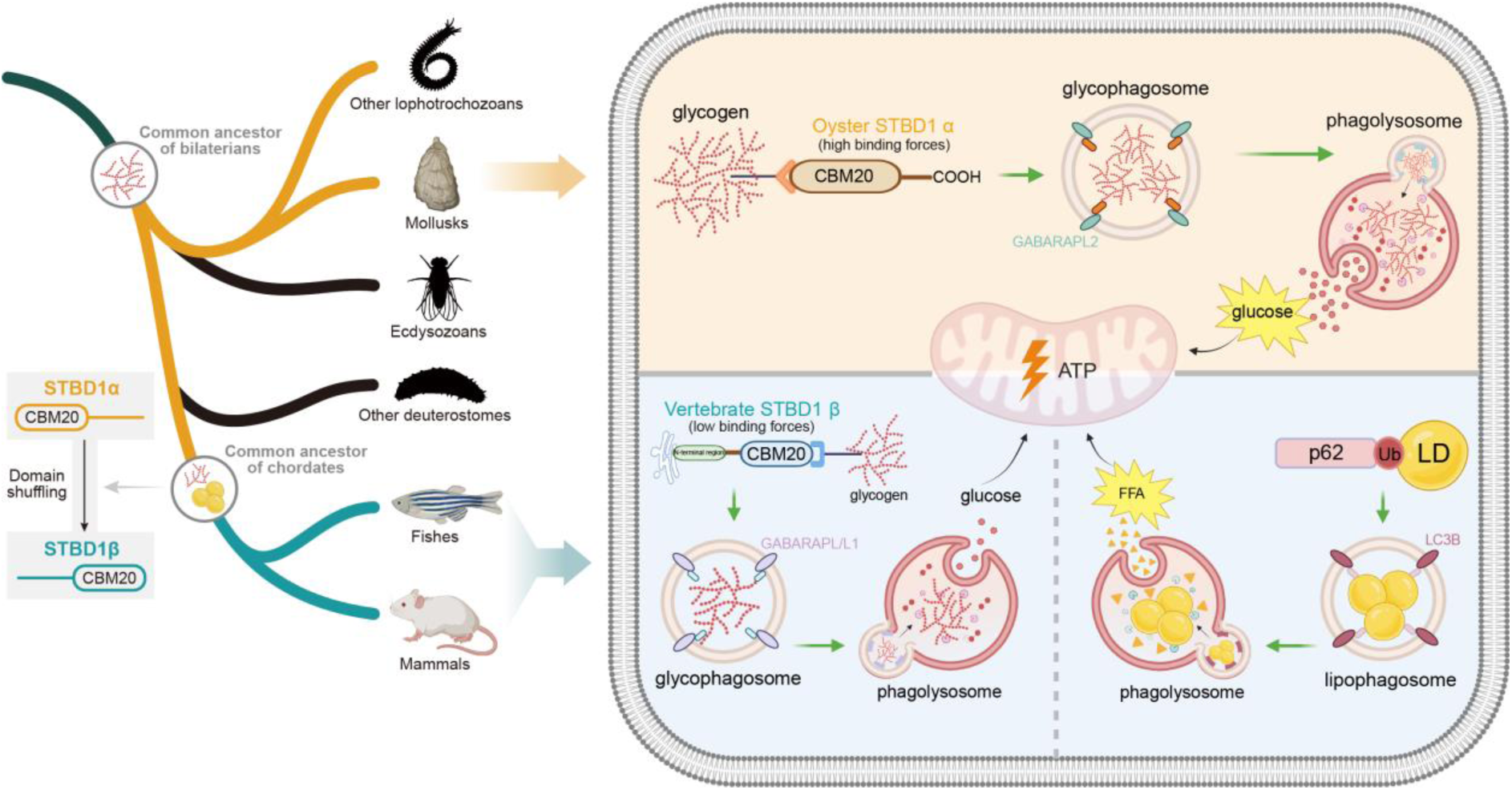
Structural evolution of STBD1 distinguishes glycophagy between oysters and vertebrates. STBD1-mediated glycophagy originated before the diversification of bilaterians. In mollusks and other lophotrochozoans, STBD1α represents the ancestral domain architecture of STBD1 proteins. Among them, oyster STBD1α binds glycogen with high affinity and subsequently anchors to GABARAPL2, facilitating glycophagic flux and providing a primary energy source. Vertebrate STBD1β proteins inherited an ancestral chordate rearrangement of the CBM20 domain to the C-terminus, resulting in lower binding affinity with glycogen compared to the N-terminal position, leading to a relatively reduced glycophagic flux through the glycophagy pathway by binding with GABARAPL or GABARAPL1. Consequently, oysters utilize glycophagy to degrade glycogen as a primary energy source, whereas vertebrates rely more strongly on lipophagy. Abbreviations: LD, lipid droplet; FFA, free fat acid.

## Materials and Methods

### Animals and treatment

All animal experiment guidelines were approved by the Institutional Animal Care and Use Committee of Ocean University of China (OUC-IACUC), with approval numbers 2020-0032- 0517 and 2023-0032-0039. Two-year-old healthy *M. gigas* individuals (*N* = 200) were collected from an oyster farm in Rongcheng (37.1°N, 122.5°E, Shandong, China) in May, and acclimated in 50 L tanks for 7 days. The oysters were subjected to a 14-day (d) period of consecutive fasting, followed by a single refeeding event with approximately 2.52 × 10^5^ cells/mL of concentrated algae fluid, and then observed for 48 hours (h). Mantle, gill, labial palp, digestive gland, adductor muscle and visceral ganglion was dissected from *N* = 3 individuals after fasting for 14 d and refeeding for 12 h, then fixed for histology. Labial palp was dissected from *N* = 6 individuals at 1 d, 3 d, 5 d, 7 d, 14 d after fasting, then again, 1 h, 3 h, 6 h, 12 h, 24 h, 48 h after refeeding for molecular work. *N* = 9 individuals were sampled at 0 d prior to fasting as a control group. Six tissues were dissected from *N* = 6 individuals for glycogen quantification (fig. S4). Labial palp was dissected from *N* = 71 randomly selected individuals for glycogen content quantification, based on the observed distribution, we selected two extreme groups (*N* = 9 each) with average dry weight glycogen content of 19.15 ± 4.05% (high glycogen) and 10.7 ± 0.45% (low glycogen) (fig. S7). All samples used for molecular work were flash frozen in liquid nitrogen and kept at -80°C.

### Glycogen content measurement and staining

Glycogen content was determined using a Glycogen Content Assay Kit following the manufacturer’s instructions (Solarbio, BC0345). Prior to measurement, the tissue samples were subjected to freeze-drying for 48h followed by pulverization into powder (0.1 g used for measurements). The cell samples were subjected to 20 rounds of sonication for 3s, at intervals of 10s. Then, 750 μL of extract solution was added to each sample, before the mixture was boiled for 20 min and centrifuged at 8,000 g for 10 min to obtain the supernatant. Finally, 240 μL of anthranone was added and boiled, before the light absorption value was determined using a Synergy™ H1 (BioTek) plate reader at an optical density (OD) of 620.

Glycogen staining was performed using a Periodic Acid Schiff (PAS) Stain Kit (Solarbio, G1281) according to the manufacturer’s instructions. Briefly, samples were fixed in 4% PFA, dehydrated in gradient alcohol, embedded in paraffin and sectioned at 5 μm thickness using a LEICA RM2016 microtome (Lecia). Sections were hydrated and oxidized with periodic acid for 5 min, stained with Schiff reagent for 15 min, and counterstained in hematoxylin for 1 min. Images were acquired with a Zeiss Axio Scope A1 microscope equipped with an AxioCam MRc5 digital camera. Positive signals were quantified using Image J (*56*) (v 2.3.0).

### Lipid staining

Lipid staining was done using Oil Red O Saturated Solution (Solarbio, G1260) according to the manufacturer’s instructions. Samples were prepared as described for glycogen staining. Sections were stained with Oil Red O solution for 10 min, differentiated in 60% isopropanol until the interstitial spaces became clear, and then counterstained in hematoxylin for 1 min for nuclear staining. Images acquisition and quantification of positive signals was performed as described for glycogen staining.

### Immunohistochemistry assay

Tissue samples were fixed in 4% PFA, dehydrated in gradient alcohol, and then embedded in paraffin and sectioned at 5 μm thickness. Sections were deparaffinized and hydrated followed by antigen retrieval in 0.01 M sodium citrate buffer (pH 6.0) at 95 °C for 10 min. Slides were rinsed by 1 × PBS for 15 min and immersed in 3% H_2_O_2_ for 10 min, then blocked with 5% BSA in PBS for 30min. Next, sections were incubated with 1:400 diluted primary antibody rabbit anti-LC3A/B (Cell Signaling Technology, 4108) overnight at 4 °C. After 1 × PBS washing for 30 min, secondary antibody HRP-conjugated goat anti-rabbit was added, and incubated for 30 min. Finally, staining was performed with 3,3-diaminobenzidine (DAB) substrate and counterstained with hematoxylin for nuclear differentiation. Negative control without primary antibody treatments were included. Images acquisition and quantification of positive signals was performed as described for glycogen staining.

### Quantitative real-time PCR

Total RNA was extracted from samples using FreeZol reagent (Vazyme, R711) according to the manufacturer’s instructions. RNA concentration and purity were assessed using a NanoDrop 2000 spectrophotometer (Thermo Fisher). Total RNA (1 μg) was reverse transcribed with the HiScript III RT SuperMix for qPCR (+gDNA wiper) (Vazyme, R323). The transcript abundances of genes of interest were measured using ChamQ SYBR qPCR Master Mix (Vazyme, R311) with a LightCycler®480 real-time PCR instrument (Roche). 10.0 μL RT-PCR reactions contained 5.0 μL 2 × SYBR Green PCR Master Mix, 0.7 μL of each forward and reverse primer (10 μM), 1.0 μL diluted cDNA, and 2.6 μL PCR-grade water. The relative expression of target genes was calculated by the 2^-ΔΔCt^ method using either EF1α (Elongation factor 1-α) or the gene combination EF1α-Arf (ADP-ribosylation factor)-GAPDH (glyceraldehyde-3-phosphate dehydrogenase) as reference (*57*). When multiple genes were used for normalization, the geometric mean of their Ct values was applied. Target gene specific primers were designed using Primer Express software (Applied Biosystems) (described in table S2). A two-tailed T test was used to assess differences in expression across different points in the time course compared to the baseline at 0 d before fasting. *P* < 0.05 was considered to represent a statistically significant difference.

### STBD1 sequence analyses

Amino acid sequences of STBD1 were downloaded from NCBI for mouse (52331), zebrafish (792854) and wild-type *M. gigas* (LOC105333128). The domain structures of these proteins were further confirmed by comparison with the InterPro database. AIM motifs were identified based on the standard sequence [TRP/PHE/TYR]-X-X-[IIE/LEU/VAL]. To compare STBD1 protein structures carrying N-terminal or C-terminal CBM20 domains, an *in silico* MgSTBD1 mutant was generated by repositioning the N-terminal CBM20 domain (aa 1-106) to the C-terminus (aa 90-194). Mouse and zebrafish STBD1 were used as comparative references containing a C-terminus CBM20 domain. STBD1 structural models were generated using the multiple threading approach LOMETS via the I-TASSER server (*58*) (https://zhanggroup.org/I-TASSER/). Models with the lowest C-score were used for molecular docking analysis described below. PyMOL (*59*) (v 2.5.5) was used for visualizing protein structures.

### Molecular docking

Molecular docking of glycogen with STBD1 proteins from mouse, zebrafish, wild-type *M. gigas*, and the *in silico M*. *gigas* STBD1 mutant was carried out using AutoDock Vina (*60*) (v 1.1.2). Protein structures were refined through the incorporation of hydrogen atoms, integration of non-polar interactions, before computation of kollman charges. The 3D structure of the ligand glycogen was obtained from the PubChem database (https://pubchem.ncbi.nlm.nih.gov/), before being optimized and energy minimized using Chem 3D (*61*), and finally hydrogen atoms were added and the torsion key was set. After pretreatment of the ligand and receptors, the parameter files were configured with appropriate grid box dimensions for STBD1 from mouse (30.75 × 24.0 × 27.75), zebrafish (29.25 × 31.5 × 26.25), *M. gigas* (25.5 × 26.25 × 28.5) and the *in silico M*. *gigas* STBD1 mutant (30.0 × 25.5 × 32.5). Running exhaustiveness was set to 10, and default settings were used for all other parameters. Protein-ligand interactions were visualized by PyMOL (*58*) (version 2.5.5) in 3D view and Ligplot (*62*) (version 2.2) in 2D view.

### Phylogenetic analysis of metazoan CBM20 domain-containing proteins

We downloaded protein datasets from 61 metazoan genomes representing 13 phyla (table S1). Proteins containing sequence homology to the CBM20 domain (Pfam: PF00686.24) were identified using hmmsearch (HMMER v.3.3.2) (*63*) employing trusted cutoffs. Domain annotation was achieved using InterProScan (v.5.69) with default parameters (*64*). Transmembrane helices were predicted using TMHMM-2.0 (*65*) online (https://services.healthtech.dtu.dk/services/TMHMM-2.0/). Whole sequences were aligned using MAFFT (*66*) (v.7.525) with the parameters “--maxiterate 1000 --localpair” (Data S1). Phylogenetic relationships were estimated using maximum likelihood with IQ-Tree (v.2.3.5) and the parameters “-B 5000 -bnni” (*67*). ModelFinder (*68*) from IQ-Tree was used to select the best-fitting model (LG+I+R6). The final phylogenetic tree was visualized and labeled using iTOL (*69*) (v.6.9.1) online (https://itol.embl.de/).

### Plasmid construction

Plasmids used for purification of recombinant STBD1 proteins were constructed in the pET32a vector (with 6 × His-tag, Biomed). The complete coding sequences of STBD1 from mouse (52331), zebrafish (792854) and wild-type *M. gigas* (LOC105333128) were PCR amplified using 2 × Phanta Max Master Mix (Vazyme, P525). The *in silico M*. *gigas* STBD1 mutant was generated by overlapping PCR based on homologous sequences of fragment-1 (nucleotides 1-318) and fragment-2 (nucleotides 319-585), which resulted in 1-106aa being rearranged to 90-194aa (Fig. 2E). The purified PCR products were subcloned into pET32a vector pretreated with BamHI and HindIII (New England Biolabs) using ClonExpress® II One Step Cloning Kit (Vazyme, C112) to generate the four pET32a-STBD1 constructs. Plasmids used for overexpression of each STBD1 protein in 293T cells were constructed in the pcDNA3.1 vector (Invitrogen). A HA-tag (TACCCATACGATGTTCCAGATTACGCT) was added at the C-terminus by PCR amplification. Then, the purified PCR products were subcloned into pcDNA3.1 vector pretreated with NheI and BamHI (New England Biolabs) to generate pcDNA3.1-HA-STBD1 expression plasmids for each of the four proteins. Plasmids used for co-IP of STBD1 from *M. gigas* with LC3A, LC3C, GABARAP and GABARAPL2 were also constructed in the pcDNA3.1 vector. The plasmid used for IP with STBD1 from *M. gigas* was constructed using the pcDNA3.1-HA expression plasmid. The LC3A, LC3C, GABARAP and GABARAPL2 expression plasmids were generated by amplifying their coding sequences with a Flag tag (GATTACAAGGACGACGATGACAAG) at N-terminus and cloning into the pcDNA3.1 vector. Primers for the construction of expression plasmids are listed in table S2. All plasmids were extracted using the EndoFree Midi Plasmid Kit (TIANGEN, DP118) and confirmed by Sanger sequencing.

### Glycogen co-sedimentation assay

Prior to the co-sedimentation assay, the four STBD1 and pET32a vector proteins were purified using a prokaryotic expression system. Briefly, each pET32a-STBD1 plasmid, additional to a pET32a blank plasmid, was transformed into the *E. coil* BL21 (DE3) strain (Vazyme, C504) and cultured in LB medium containing ampicillin. The cultured bacteria were proliferated to OD = 0.6 and induced by 1 mM isopropyl β-D-thiogalactoside for 4 h at 37 °C. The bacteria were centrifugated and resuspended in a binding buffer (0.1 M N_2_HPO_4_, 0.01 M NaH_2_PO_4_, 0.5 M NaCl, 8 M urea and 25 mM imidazole, pH = 7.4) before ultrasonic cracking. Lysed protein supernatant was loaded on Ni-NTA agarose gel and eluted by 250 mM imidazole. Finally, proteins were refolded and further purified in ultrafiltration tubes containing dialysate buffer (2 mM Reduced Glutathione, 0.2 mM Oxidized Glutathione, 0.1M Glycine, 5% Glycerine and gradient urea [6 M-4 M-2 M-1 M-0 M]).

The co-sedimentation assay was performed as described previously (*70*). 360 μL of 0.55 mg/ml purified STBD1 proteins were incubated in 1440 μL co-sedimentation buffer (10 mM Tris-HCl, pH 7.5, 1 mM 2-mercaptoethanol, 1 mM PMSF, 1 mM EDTA and 1 mM protease inhibitor cocktail (Beyotime, P1005)) with or without 600 μL of 17.6% glycogen from cattle liver (Sigma, G0885) overnight at 4 °C. Then, 4 mL extra buffer was added, and the solution layered over 6.0 ml of 0.25 M sucrose in a 13.2 ml centrifuge tube (Beckman, 344059). After centrifugation at 100,000 g for 90 min, the supernatant and pellet were collected for Western blotting. The fraction before centrifugation was used as input. Pet32a vector protein was used as a negative control and treated in the same way.

### Overexpression of STBD1 protein in 293T cells and EBSS treatment

293T cells were cultured in DMEM (Gibco) supplemented with 10% FBS (Hyclone) and 1 × penicillin/streptomycin (Sangon) with 5% CO_2_ at 37 °C. Before transfection, 1 × 10^6^ cells per well were seeded into 6-well dishes (Corning), allowing for cellular proliferation until reaching 50-70% confluence. Then, 1 μg of each pcDNA3.1-HA-STBD1 expression plasmid was transfected into these cells using Lipofectamine™3000 (Invitrogen, L3000015) according to the manufacturer’s instructions. Cells were cultured for 24 h, followed by treatment with or without the autophagy inducer Earle’s balanced salt solution (EBSS) for 6 h. After transfection, the protein was extracted by lysis buffer (Beyotime, P0013) for Western blotting and the glycogen content within the cells determined.

### Dual-fluorescence in situ hybridization

The labial palp was used to analyze co-localization of mRNA from *STBD1*, *LC3A*, *LC3C*, *GABARAP* and *GABARAPL2* of *M. gigas*. Briefly, the samples were fixed in 4% PFA at 4°C, dehydrated in gradient alcohol, then embedded in paraffin and sectioned at 5 μm thickness. Digoxigenin-labeled *STBD1* and biotin-labeled *LC3A*, *LC3C*, *GABARAP* and *GABARAPL2* probes were transcribed *in vitro* using an RNA labeling kit (Roche, 10999644001) from the PCR products containing *STBD1*(nucleotides 7-418), *LC3A* (nucleotides 68-358), *LC3C* (nucleotides 57-320), *GABARAP* (nucleotides 71-350) and *GABARAPL2* (nucleotides 42-302) fragments. Primers for synthesizing RNA probes are listed in table S2. Sections were deparaffinized, hydrated, and then treated with 4 μg/mL proteinase K (Roche) before pre-hybridization in 66% formamide and incubation with a mixture of the *STBD1* probe with either *LC3A*, *LC3C*, *GABARAP* or *GABARAPL2* probes for 24 h at 65 °C. Slides were rinsed in maleic acid buffer and blocked with 1% blocking reagent (Roche). Subsequently, Alexa-488 conjugated anti-digoxigenin (Jackson ImmunoResearch, 200-542-156) and Alexa-647 conjugated anti-biotin (Jackson ImmunoResearch, 200-602-211) diluted 1:500 with blocking reagent were added and incubated overnight at 4 °C. Nuclear differentiation was performed using DAPI staining solution according to the manufacturer’s instructions (Beyotime, C1006). All the images were acquired with a Lecia TCS P8 laser confocal microscope.

### Co-immunoprecipitation assay

Protein interactions of *M. gigas* STBD1 with LC3A, LC3C, GABARAP or GABARAPL2 were analyzed using co-immunoprecipitation (co-IP) assays. Prior to co-IP, 293T cells were seeded at 5 × 10^6^ cells per 10 cm dish and transfected at 50-70% confluency with the above described pcDNA3.1-HA-STBD1 and either pcDNA3.1-Flag-LC3A, pcDNA3.1-Flag-LC3C, pcDNA3.1-Flag-GABARAP or pcDNA3.1-Flag-GABARAPL2 plasmids using Lipofectamine™3000 according to the manufacturer’s instructions. Transfections were cultured for 48 h at 37 °C, with 5% CO_2_, before harvest. Cells were lysed using IP lysis buffer (Beyotime, P0013), before co-IP was performed using an immunoprecipitation kit with Flag-tagged protein (Beyotime, P2202). After protein quantification using a BCA protein quantification kit (Vazyme, E112), 5% lysates were used as input, and the remaining samples incubated with Flag-tagged agarose beads overnight at 4 °C. The beads were washed 3 times with lysis buffer and the protein products eluted for Western blotting.

### Western blotting

Protein samples were denatured by boiling in 2 × SDS-PAGE loading buffer (EpiZyme, LT101) at 100 °C for 5 min. Then, proteins were separated by 12% or 15% sodium dodecyl sulfate-polyacrylamide gel electrophoresis and transferred to PVDF membranes (Millipore, America). Membranes were blocked with 5% skimmed milk dissolved by TBST buffer at 37 ℃ for 2 h. For the fasting and refeeding samples, anti-STBD1 rabbit pAb (AntibodySystem, PHB57101) or anti-p62 rabbit pAb (AntibodySystem, PHG56101) were added (1:2000 dilutions) and incubated overnight at 4 ℃. Other primary antibodies were added with 1:1500 dilution and incubated overnight at 4 ℃: anti-His mouse mAb (Beyotime, AF5060) for the glycogen co-sedimentation assay; anti-HA rabbit mAb (Beyotime, AG8057) for STBD1 overexpression experiments; anti-HA rabbit mAb and anti-Flag rabbit mAb (Beyotime, AG8057) for co-IP assays. Next, the membranes were washed 5 times with TBST and incubated with secondary antibody horseradish peroxidase (HRP)-conjugated goat anti-mouse IgG (Beyotime, A0216) or goat anti-rabbit IgG (Beyotime, A0208) diluted 1:1000 by TBST for 1 h at 37 °C. β-actin mouse mAb (Beyotime, AF0003) was used as a control for the fasting and refeeding samples. GAPDH mouse mAb (Beyotime, AF5009) was used as a control for the STBD1 overexpression experiment and co-IP assays. Finally, the blots were measured with SuperPico ECL chemiluminescence kit (Vazyme, E422) and visualized using the GE ImageQuant LAS4000mini system.

## Data availability

Support information—This article contains supporting information

## Acknowledgments

This work was supported by grants from the National Natural Science Foundation of China (32341060 and 42276112), the Key Research and Development Program of Shandong Province (2021ZLGX03), the National Key Research and Development Program of China (2022YFD2400300), the Fundamental Research Funds for the Central Universities (202261032), and the earmarked fund for the Agriculture Research System of China (CARS- 49). DJM received support from an Institute Strategic Programme award (BBS/E/RL/230001C) from the Biotechnology and Biological Sciences Research Council to The Roslin Institute. We acknowledge the support of the High-Performance Biological Supercomputing Center at the Ocean University of China, and the Marine Biodiversity and Evolution Research Institute’s Instrument and Equipment Sharing Platform for providing the high-speed centrifuge (XPN- 100) for this research.

## Author contributions

SL conceived and designed the study. LR, YT, and SZ collected the samples and executed the experiments. LR, YB, CS, SL and DJM analyzed the data. LR and YB drafted the manuscript, and SL and DJM revised the manuscript. SL and QL supervised the study and provided resources. All authors have read and approved the final version of the manuscript.

## Data and materials availability

All data are available in the main text or the supplementary materials.

