## Supplementary Figures 1-15 for "Glycophagy is an ancient bilaterian pathway supporting metabolic adaptation through STBD1 structural evolution"

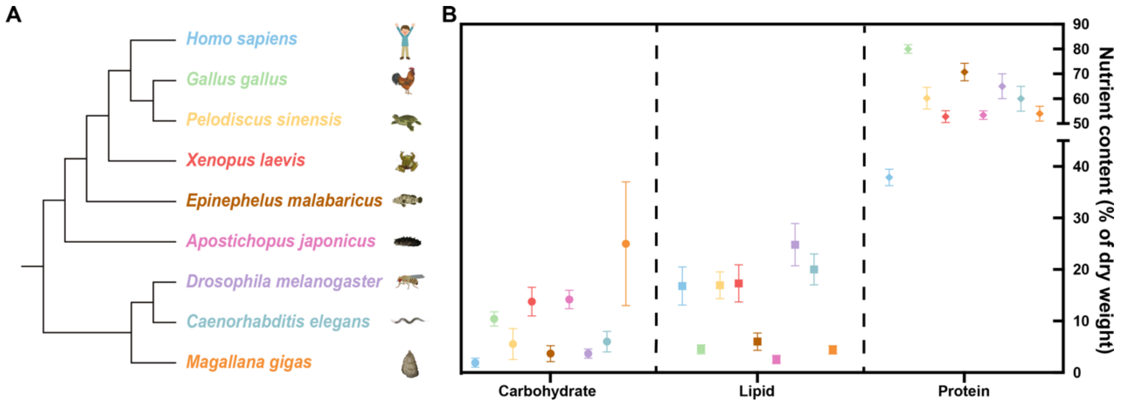


**Fig. S1. Carbohydrate, lipid and protein storage across the Metazoa. (A)** Phylogenetic tree for representative species. **(B)** Dry weight composition of carbohydrates, lipids, and proteins in *Homo sapiens* (adult man) was 1.91 ± 0.87%, 16.8 ± 3.7%, and 37.88 ± 1.6% (*1-3*); *Gallus gallus* was 10.4 ± 1.39%, 4.50 ± 0.57% , and 79.98 ± 1.69% (*4*); *Pelodiscus sinensis* was 5.52 ± 3%, 16.94 ± 2.6%, and 60.22 ± 4.37% (*5*); *Xenopus laevis* (male) was 13.7 ± 2.77%, 17.3 ± 3.6%, and 52.8 ± 2.4% (*6*); *Epinephelus malabaricus* was 3.67 ± 1.55%, 6.00 ± 1.67%, and 70.74 ± 2.52% (*7*); *Apostichopus japonicus* was 14.17 ± 1.79%, 2.54 ± 0.21%, and 53.42 ± 1.22% (*8*); *Drosophila melanogaster* was 3.69 ± 0.9%, 24.8 ± 4.1%, and 65 ± 5% (*9*, *10*); *Caenorhabditis elegans* was 6 ± 2%, 20 ± 3%, and 60 ± 5% (*11*), and *Magallana gigas* was 25 ± 12%, 4.4 ± 0.8%, and 54 ± 3% (*12*).

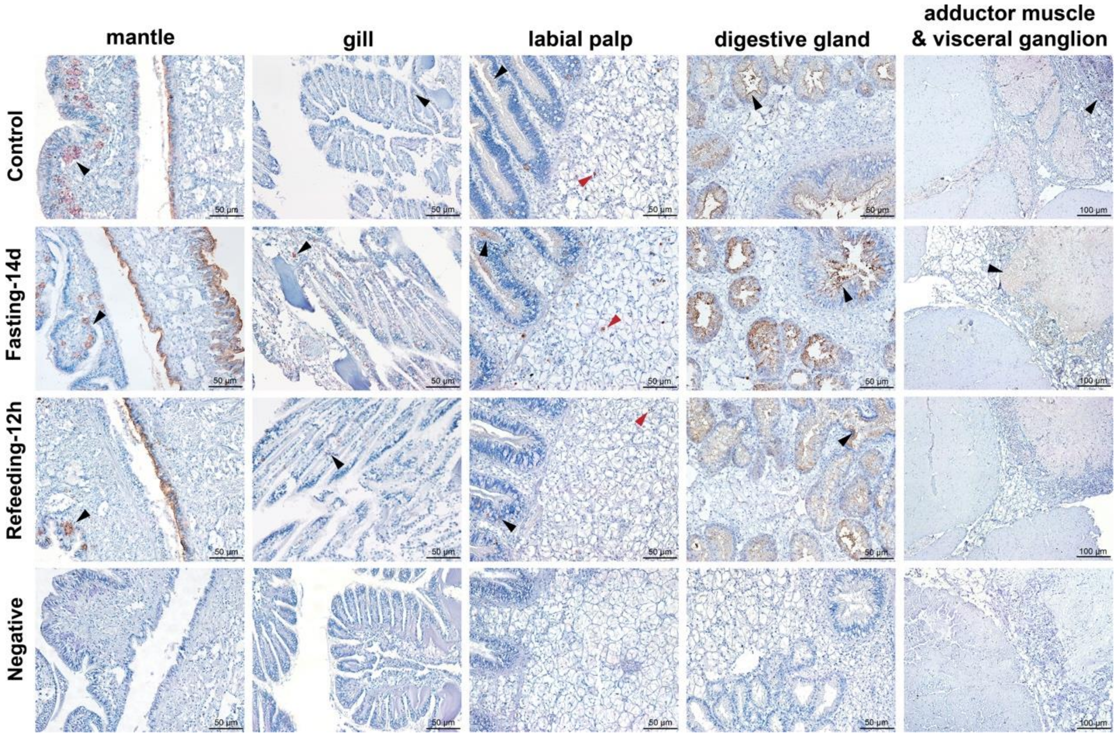


**Fig. S2. Immunohistochemistry of the autophagy marker LC3 in mantle, gill, labial palp, digestive gland, adductor muscle and visceral ganglion for control, fasting 14d and refeeding 12h groups.** Arrow indicates dense areas of LC3 positive signals such as the fold of mantle, gill filament, CE and VCT (red) cells of labial palp, digestive diverticula and visceral ganglion. No LC3 primary antibody was added in negative controls. Scale bars represent 50 μm or 100 μm.

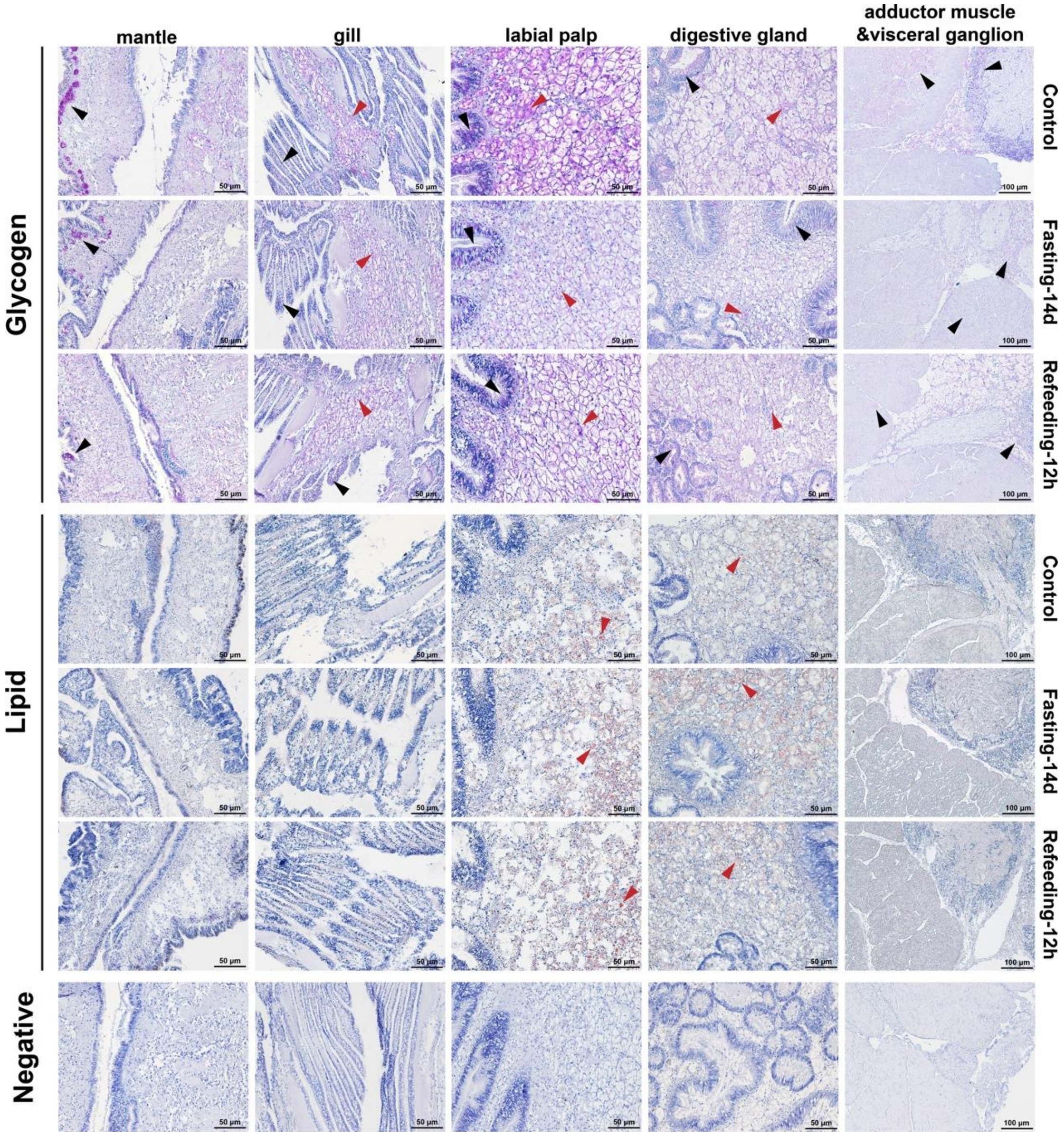


**Fig. S3. Glycogen staining and lipid staining in mantle, gill, labial palp, gonad, digestive gland, adductor muscle and visceral ganglion after fasting 14d and refeeding 12h.** Black arrows indicate dense areas of glycogen (pink) and lipid (red) signals such as the fold of mantle, gill filament, CE of labial palp, digestive diverticula, adductor muscle and visceral ganglion. Red arrows highlighted signals in the VCT cells inside each tissue. No glycogen or lipid stains were added in the negative group. Scale bars represent 50 μm or 100 μm.

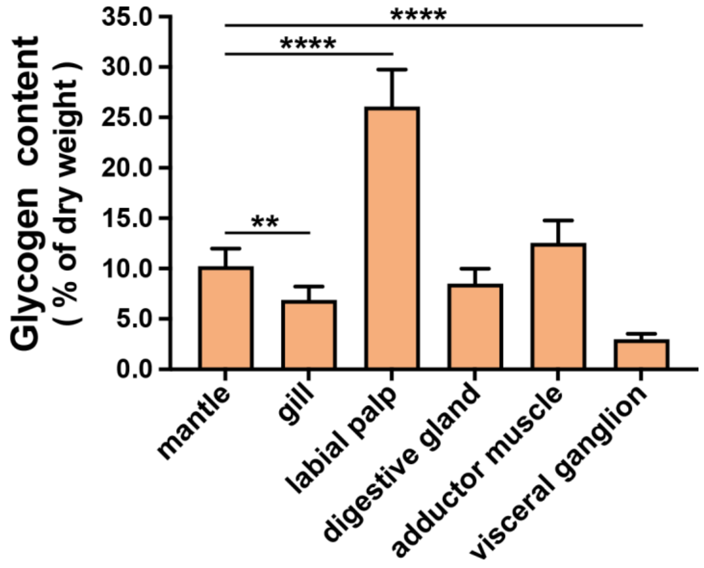


**Fig. S4. Glycogen content in the mantle, gill, labial palp, digestive gland, adductor muscle and visceral ganglion.** $N = 6$. Data represents mean ± SD. $^{**}P < 0.01$.

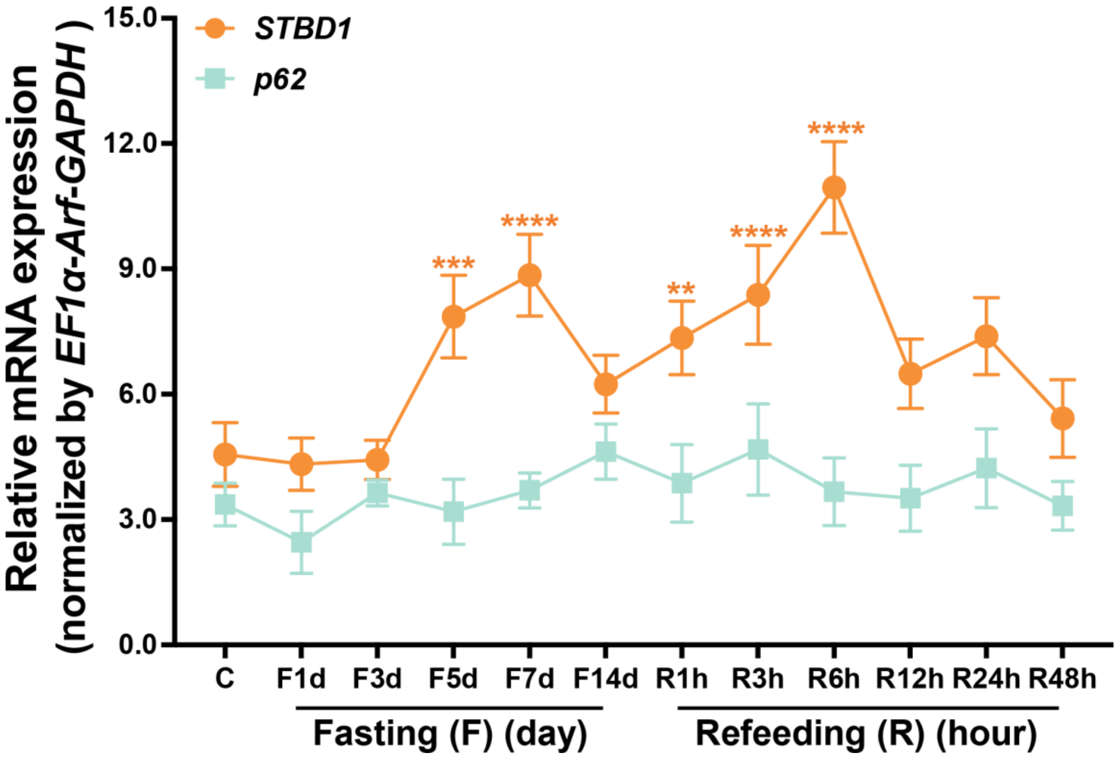


**Fig. S5. mRNA levels of STBD1 and p62 in the labial palp of oysters after fasting for 14 d and refeeding for 48 h, normalized by *EF1α-Arf-GAPDH*.** $N = 6$. Data represents mean ± SD.**$P < 0.01$, ***$P < 0.001$, ****$P < 0.0001$. F, Fasting; R, Refeeding; d, day; h, hour.

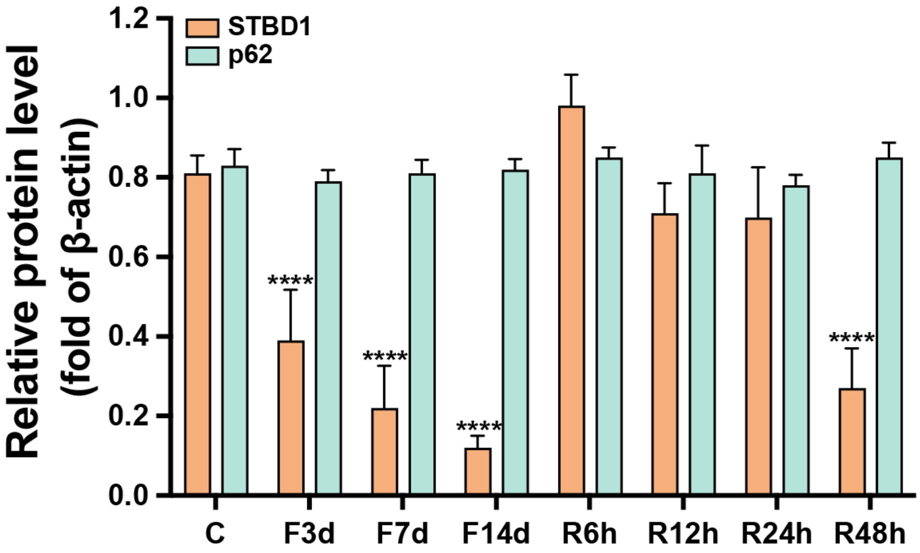


**Fig. S6. Estimated protein levels of STBD1 and p62 during fasting and refeeding, which was quantified and normalized to β-actin.** Data represent mean ± SD. ****$P < 0.0001$.

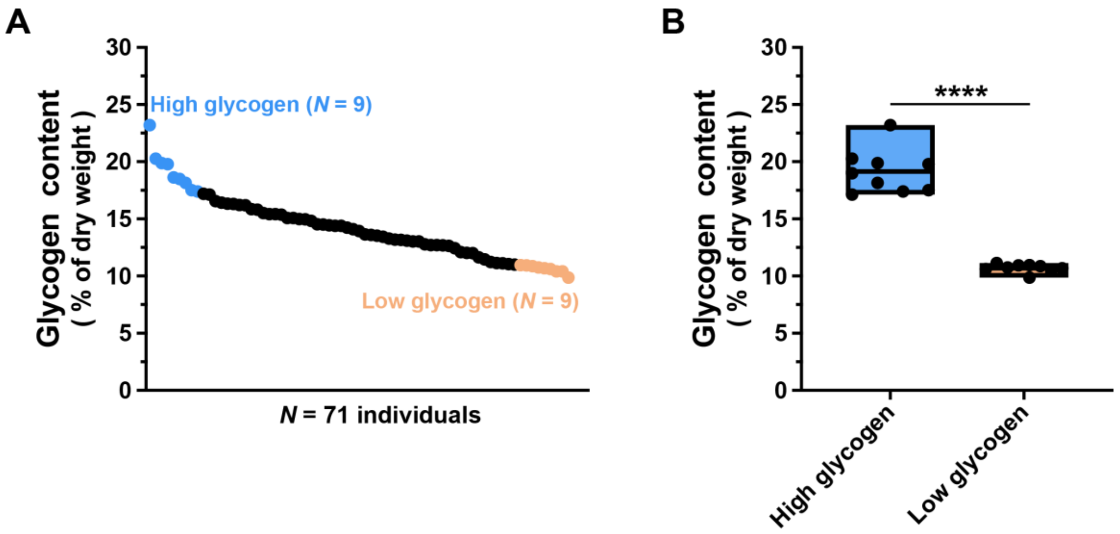


**Fig. S7. Glycogen content in high glycogen and low glycogen groups. (A)** Glycogen content in the labial palp dissected from 71 oyster individuals. Blue and pink points represent oyster samples with high and low glycogen, respectively. **(B)** The highest (average 19.15%) and lowest (average 10.7%) glycogen contents of 9 individuals sampled from 71 individuals, defining the high and low glycogen groups, respectively. Data represent mean. ****$P < 0.0001$.

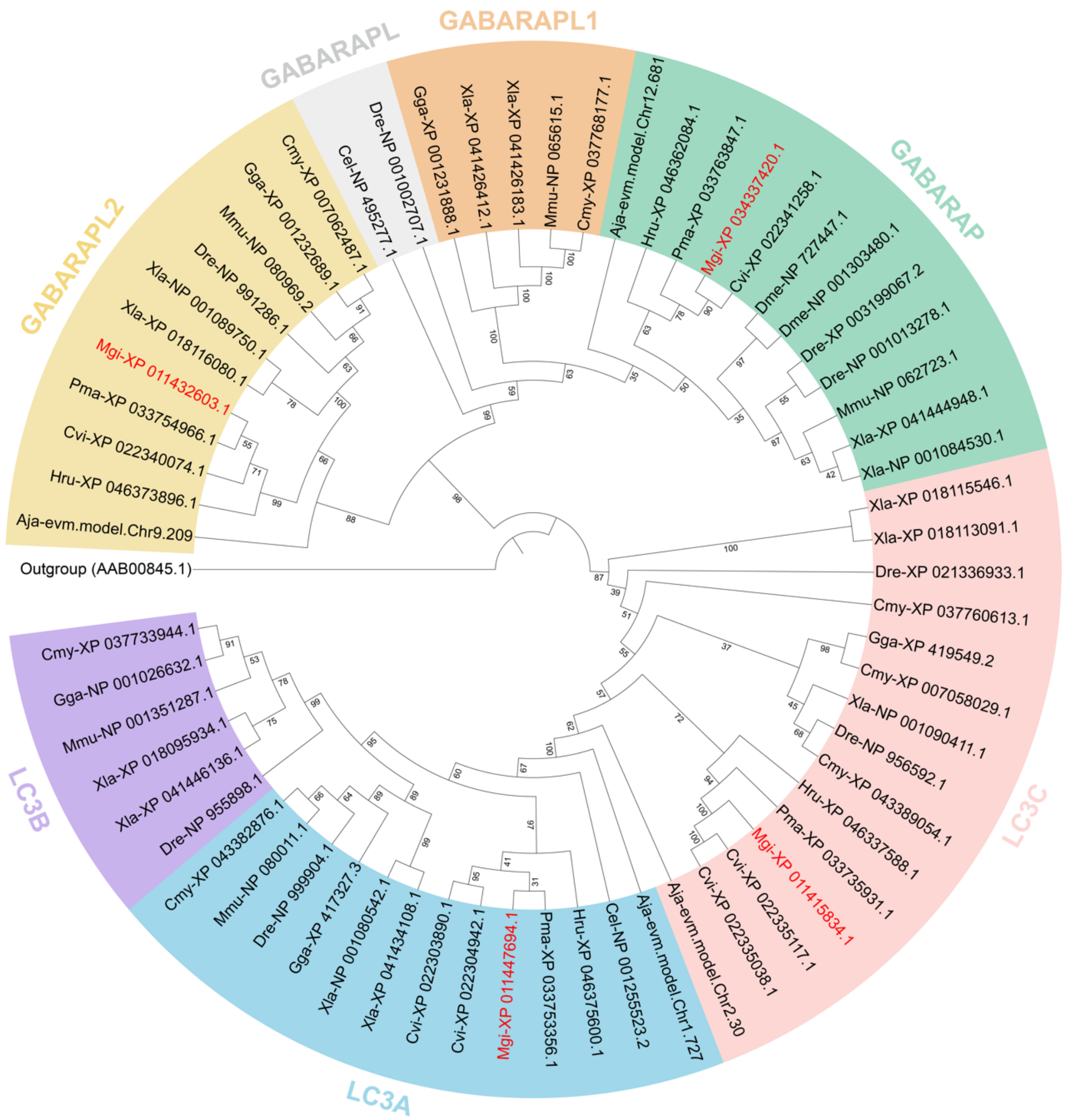


**Fig. S8. Phylogenetic tree of Atg8 family in *M. gigas* in relation to the other metazoans.** *M. gigas* sequences are highlighted in red letters. The corresponding abbreviations of species names are shown in Table S1. The alpha-cyclodextrin glycosyltransferase from *Thermoanaerobacterium thermosulfurigenes* (GenBank accession: AAB00845.1) was used as an outgroup.

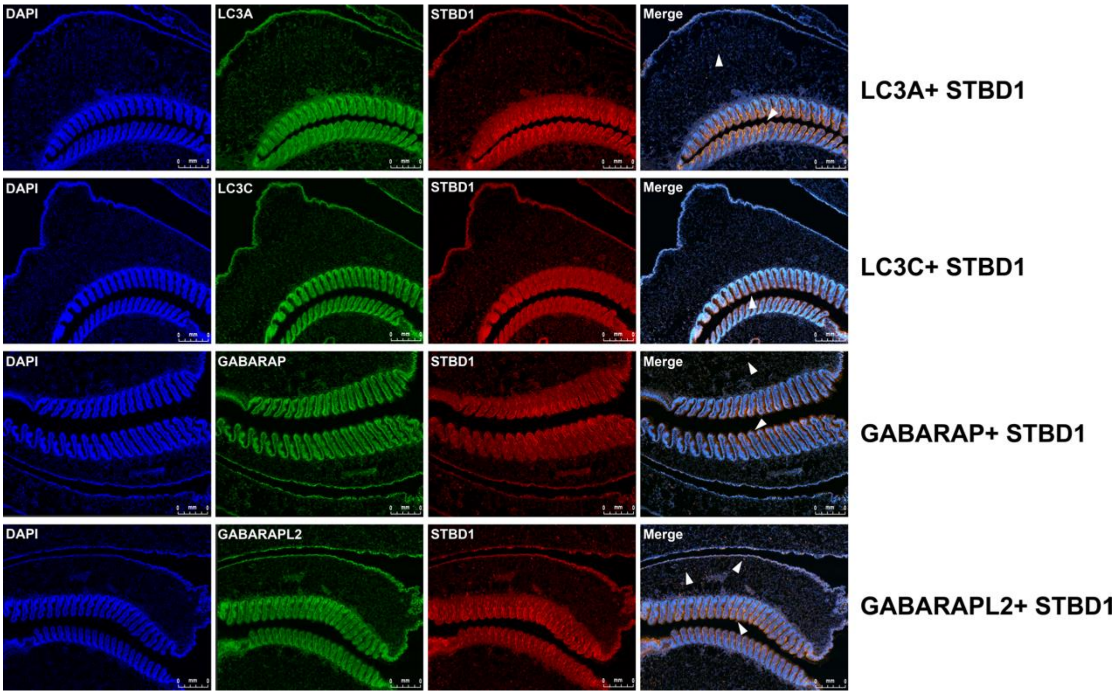


**Fig. S9. Fluorescence *in situ* RNA hybridization detects co-localization of transcripts for *STBD1* and either *LC3A*, *LC3C*, *GABARAP* or *GABARAPL2* in the labial palp of oysters.** Signals from nucleus (blue), *LC3A*, *LC3C*, *GABARAP* and *GABARAPL2* (green), *STBD1* (red), and merge. *STBD1* co-localized with *LC3A*, *LC3C*, *GABARAP* or *GABARAPL2* signals were indicated with white arrows.

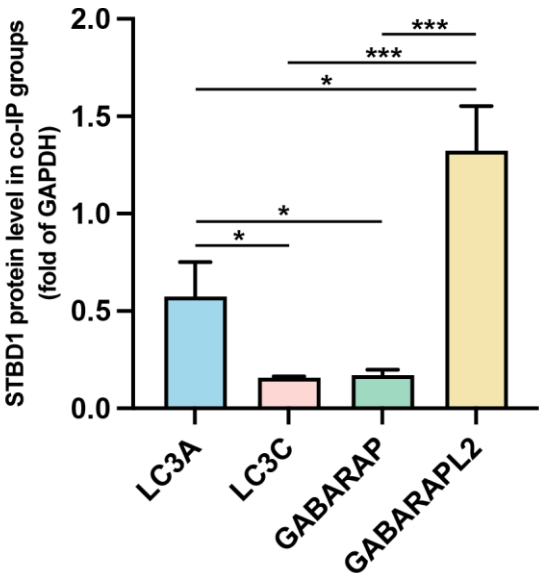


**Fig. S10. Estimated protein levels of STBD1 in complex with either LC3A, LC3C, GABARAP or GABARAPL2 respectively, normalized to GAPDH.** Data represents mean ± SD. **P* < 0.05, ****P* < 0.001.

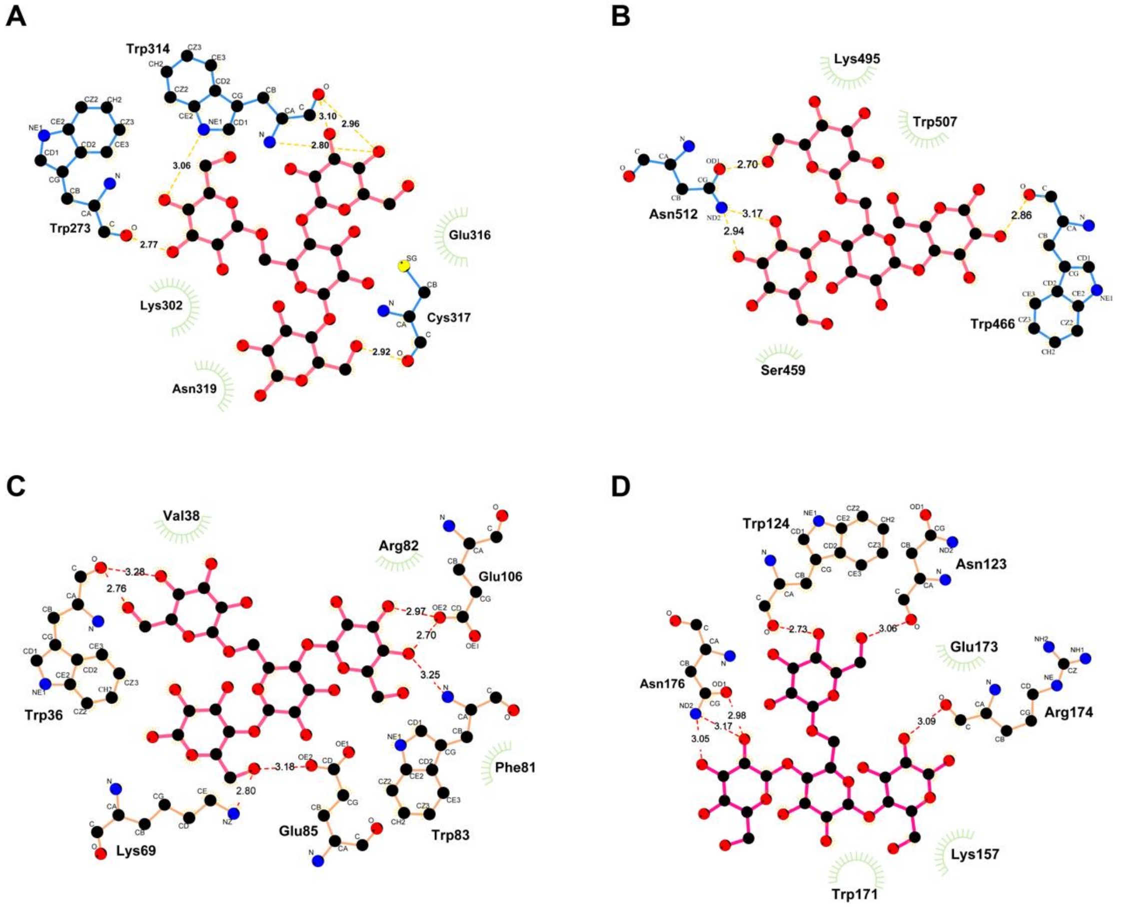


**Fig. S11. Two dimensional views of docking models of glycogen with STBD1 from mouse (A), zebrafish (B), *M. gigas* (C), and a *M. gigas* STBD1 in silico mutant with CBM20 moved to the C-terminus (D).** Hydrophobic residues are indicated by green radiations.

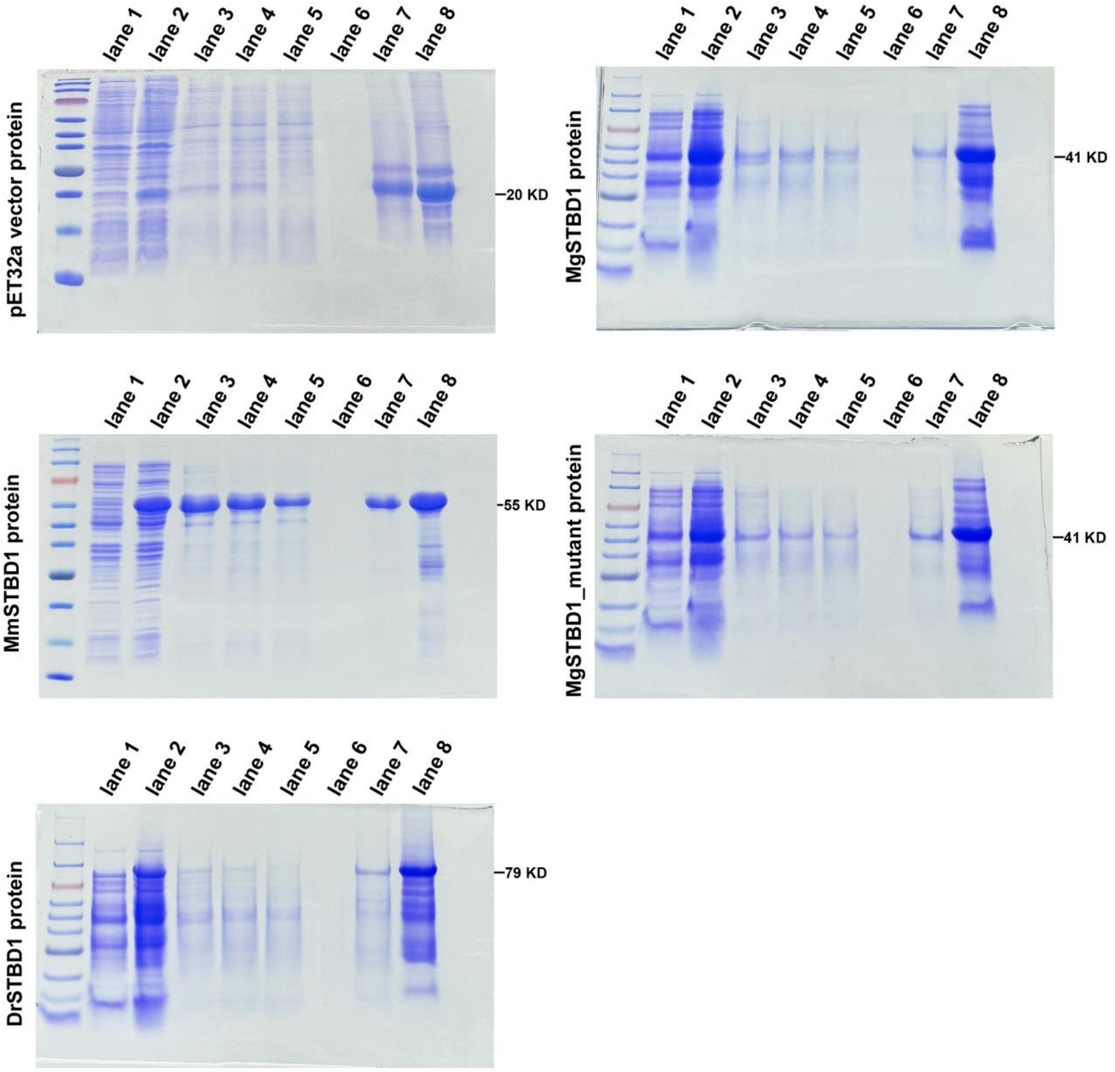


**Fig. S12. SDS-PAGE was used to verify the molecular weights of pET32a vector recombinant proteins for STBD1 from mouse, zebrafish, wide-type *M. gigas* and a mutant *M. gigas* STBD1 with CBM20 moved to the C-terminus.** Lane 1: prior to induction; lane 2: post induction; lane 3: bacterial lysate; lane 4: post-lysis supernatant; lane 5: flow-through fraction; lane 6: washing buffer; lane 7: elution buffer; lane 8: concentration buffer. Abbreviations used: Mm, *Mus musculus*; Dr, *Danio rerio*; Mg, *Magallana gigas*.

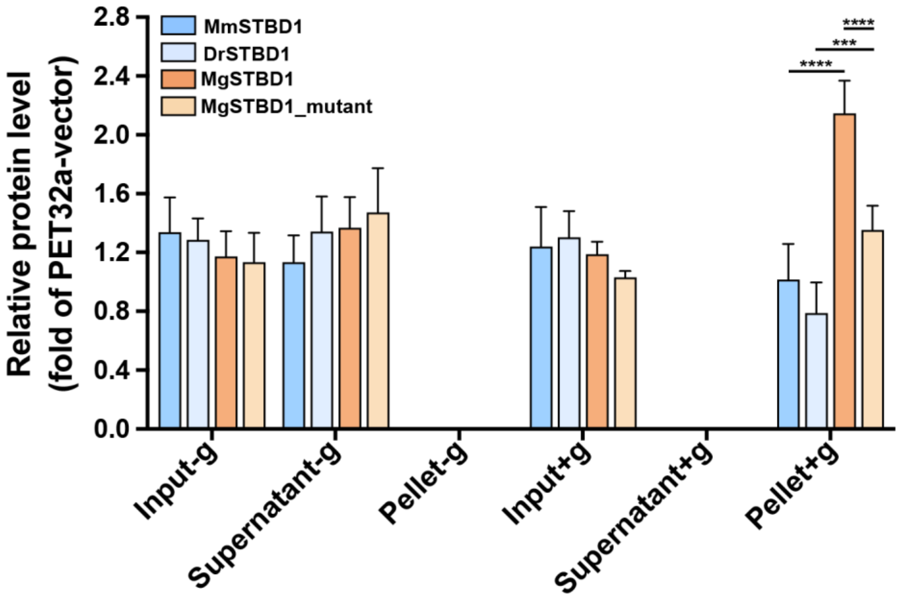


**Fig. S13. Estimated abundance of binding between glycogen and STBD1 from mouse, zebrafish, wide-type *M. gigas* and a mutant *M. gigas* STBD1 with CBM20 moved to the C-terminus, normalized to PET32a vector protein level.** Data represents mean ± SD. ***$P$ < 0.001, ****$P$ < 0.0001.

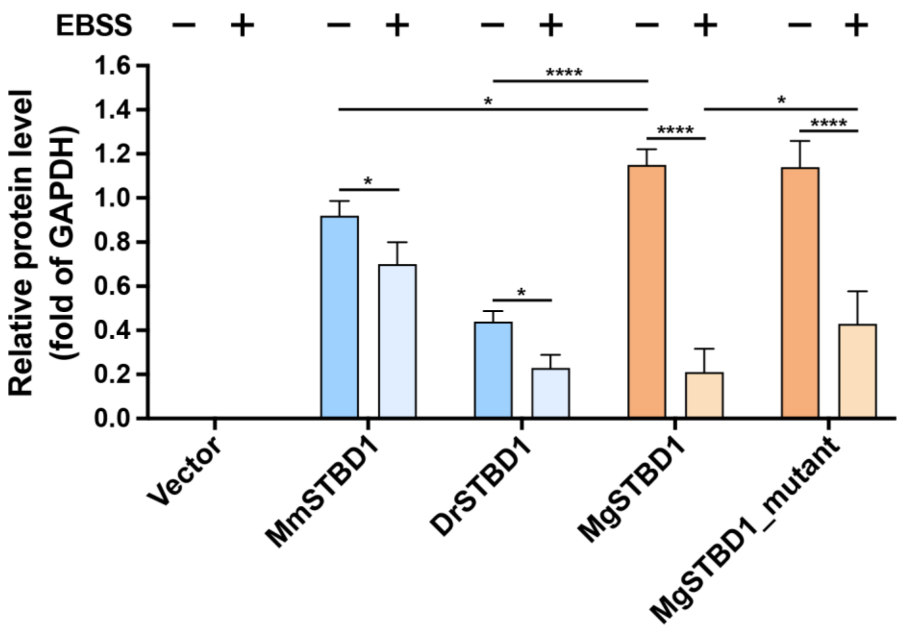


**Fig. S14. Estimated abundance of protein levels for STBD1 from mouse, zebrafish, wide-type *M. gigas* and a mutant *M. gigas* STBD1 with CBM20 moved to the C-terminus, overexpressed in 293T cells with or without EBSS treatments, normalized to GAPDH levels.** Data represents mean ± SD. *$P < 0.05$, ****$P < 0.0001$.

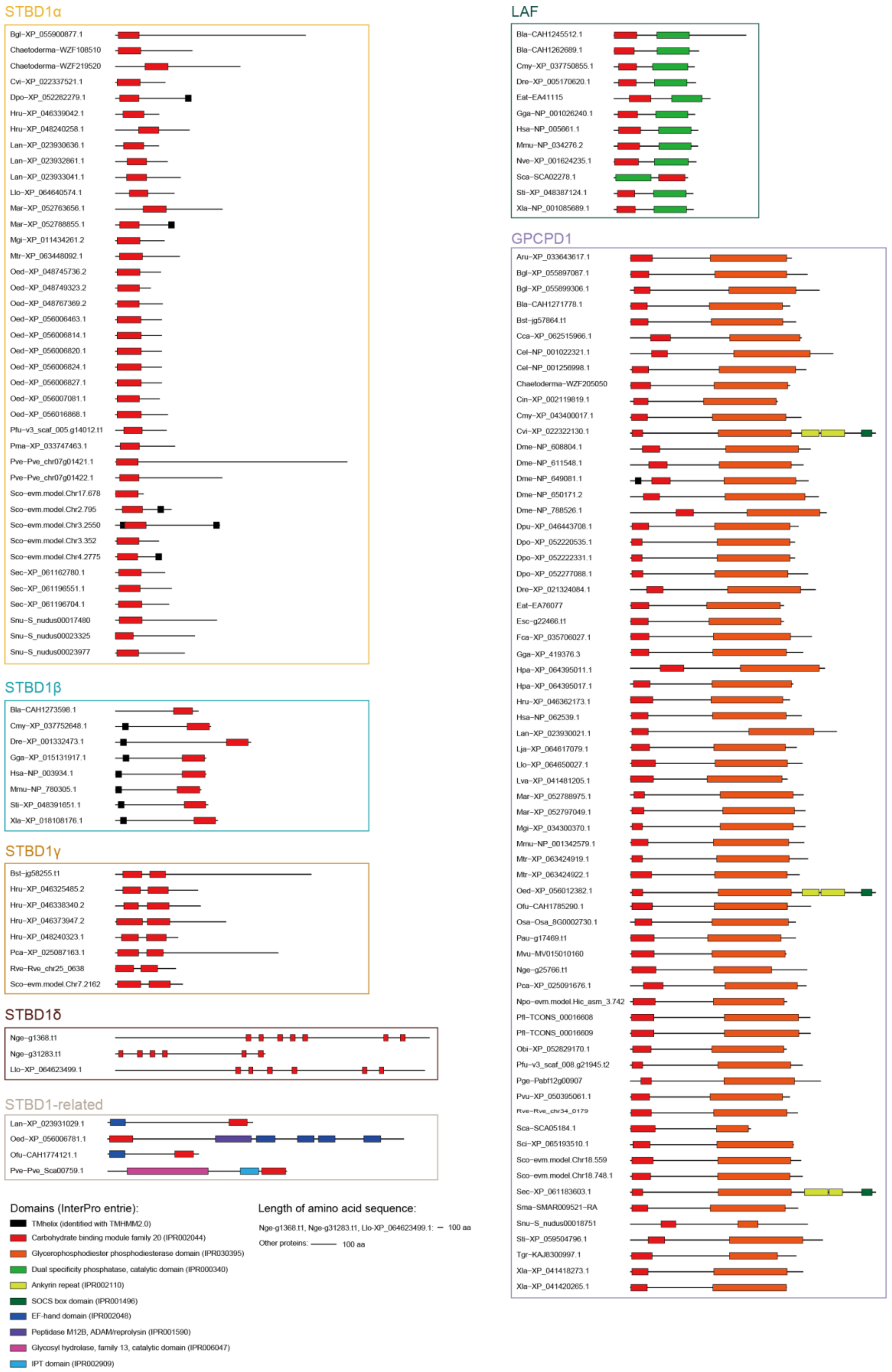


**Fig. S15. Domain arrangements of CBM20 containing proteins.** The proteins containing CBM20 domain(s) were classified into seven types based on domain architectures. Briefly, among all identified proteins, those with CBM20 and dual specificity phosphatase domains were considered as LAF; those with CBM20 and glycerophosphodiester phosphodiesterase domains as GPCPD1; those with a single N-terminal CBM20 domain as STBD1α; those with one C-terminal CBM20 domain as STBD1β; those with two N-terminal CBM20 domains as STBD1γ; those with more than two N-terminal CBM20 domains as STBD1δ; and those with other domains such as EF-hand, peptidase M12B, IPT or glycosyl hydrolase family 13 as STBD1-related proteins. The corresponding abbreviations of species names are shown in table S1. Our classification of STBD1, LAF and GPCD1 proteins was further supported by their monophyly in our phylogenetic analysis (Fig. 4).
